# Non-cognate molecular interactions impact bacterial growth

**DOI:** 10.1101/2021.10.20.465141

**Authors:** Claudia Igler, Bryan Wu, Claire Fourcade, Bor Kavčič, Torsten Waldminghaus, Florian M. Pauler, Balaji Santhanam, Gašper Tkačik, Călin C. Guet

## Abstract

In gene regulatory networks, appropriate gene expression is often determined through high-affinity binding of transcription factors (TFs) to their cognate DNA sequences. However, the large genomic background likely contains many DNA sequences similar to the TF target motifs, potentially allowing for ample low-affinity non-cognate TF binding. Whether and how non-cognate TF binding impacts cellular function and growth remains unclear. Here, we find that increased expression of native and non-native TFs in *Escherichia coli* and *Salmonella enterica* can significantly inhibit population growth across different growth conditions. This effect depends upon low-affinity TF binding to many DNA sequences, TF cooperativity, and TF concentration. Non-cognate TF binding to DNA can thus severely impact fitness, giving rise to fundamental biophysical and physiological constraints on the evolution of gene regulatory networks and also their synthetic design.

## Introduction

All levels of biology depend upon timely recognition and interactions between cognate biomolecules (Box 1A). The importance of molecular encounters between intended targets is highlighted by the intricate mechanisms that ensure cognate – and suppress non-cognate – interactions, a classic example being that of kinetic proofreading in the loading of amino acids onto tRNAs (Hopfield, 1974). Gene expression is a tightly regulated process that is modulated by transcription factors (TFs) binding to their respective cognate operator sites, thereby determining the right time and place for gene expression. This binding is usually with high affinity, although functional binding with low affinity to “weak” binding sites has also been recognized, especially in eukaryotes (Kribelbauer *et al*., 2019; Shahein *et al*., 2022). Additionally, non-cognate low-affinity interactions can occur at many other DNA sequences, as TFs will typically have some propensity to bind even sequences far away from their consensus (Kribelbauer *et al*., 2019). Given the typical genome size of several million base pairs, such sequences are likely to be encountered many times within a genome. These non-cognate interactions are not necessarily detrimental; for example, non-cognate DNA binding was found to speed up the TF search process by sliding along the DNA in a process called “facilitated diffusion” (Elf *et al*., 2007; Mirny *et al*., 2009), where conformational changes upon encountering the target sequence result in cognate binding (Mirny *et al*., 2009). In prokaryotes, many transcriptional regulators appear to be bound to non-cognate DNA most of the time (Elf *et al*., 2007; Flyvbjerg *et al*., 2006; Stracy *et al*., 2021) and this state could play an important role in gene regulation (Bakk & Metzler, 2004b; Kao-Huang *et al*., 1977; von Hippel *et al*., 1974).

Non-cognate TF-DNA interactions are often referred to as non-specific as they are likely very weak and transient (Bakk & Metzler, 2004a; Elf *et al*., 2007). It is important to realize, however, that specificity is a biophysical property of a TF, describing – depending on the field – either the distribution of the TF’s residence time when bound to different sequences, or the TF’s binding mode and structural conformation upon interacting with the DNA (Jonge *et al*., 2022) (Box 1). Whether any such binding is biologically functional and has been selected for (cognate) or spurious (non-cognate) is a separate question. In this manuscript, we will focus on binding with high or low affinity at either cognate or non-cognate DNA sites.

**Box 1.**
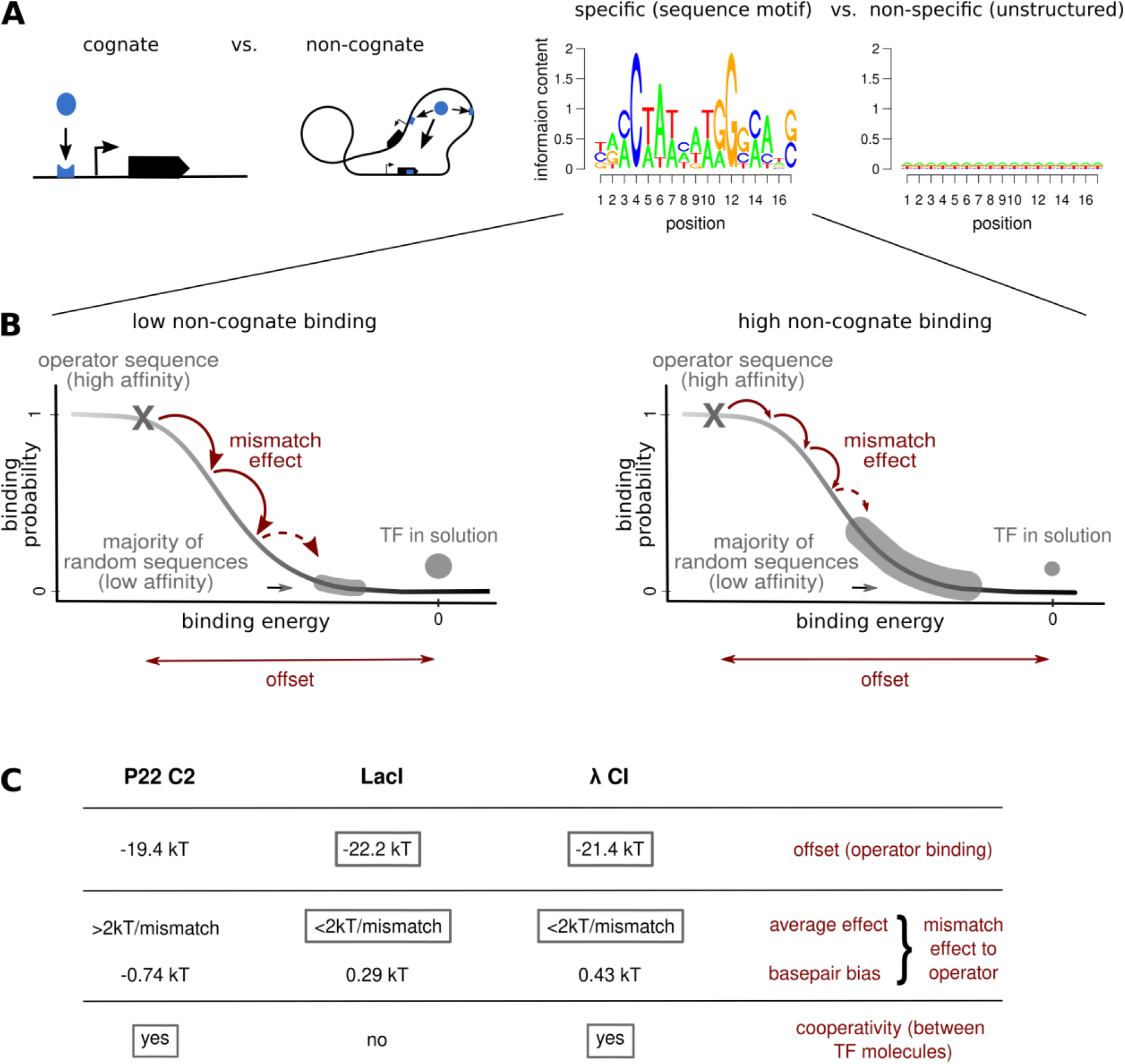
The biophysics and molecular biology of TF crosstalk. **(A)** In order to elicit an appropriate function, TFs (blue ellipsoid) have to recognize their cognate DNA sites known as operators (blue trough) among a large background of non-cognate sites (blue rectangle), where binding is non-functional (i.e., off-target) and potentially interferes with cellular programs. Binding of TFs to cognate or non-cognate sites can occur by recognizing a TF-specific sequence motif on the DNA (with binding affinity ranging from high to low, depending on the similarity of the target site with the TF’s consensus sequence (Gerland et al., 2002)) or through generic, inherently low-affinity interactions with any DNA sequence. **(B)** Schematics of binding probability curves for TFs with lower (e.g. P22 C2, left panel) or higher (e.g. λ CI, right panel) likelihood to bind non-cognate DNA targets are shown. Binding with high affinity usually occurs to a TF-specific DNA motif and is determined by three main factors: (i) the offset gives the binding energy (lower energy = stronger binding) to a single operator sequence (usually the strongest wild-type operator sequence, gray cross) relative to the unbound state (TF in solution; gray dot above the sigmoid). Random DNA sequences are usually bound with low affinity and are located at the lower end of the sigmoid (gray curved box) with energy values approaching that of the TF in solution. The number of TFs being bound to random sequences or free in solution are indicated by the size of the gray box and dot. (ii) Basepair mismatches to the wild-type operator sequence incur an energy penalty and increase the binding energy (i.e. weaken the binding; shown by the red arrows for each mismatch) based on motif-dependent basepair binding energy mismatches and overall basepair bias (given here as average AT preference – average GC preference). Sequences with a higher number of mismatches to the operator sequence will be more abundant than ones with few mismatches, which is symbolized by the shading of the binding sigmoid. Hence, for a TF with large negative offset, such as λ CI, which is aided by (iii) high cooperativity, and an energy matrix characterized by small mismatch effects, many random sequences will not be far from or even on the rise of the sigmoid, making higher occupancy likely (see C). **(C)** Table compares these three factors (offset, basepair mismatches and cooperativity) for three well-characterized TFs, showing that all of them are fulfilled for λ CI, but only partially for P22 C2 and LacI (Barnes et al., 2019; Hilchey et al., 1997; Igler et al., 2018; Vilar & Saiz, 2005), making λ CI an obvious candidate for substantial non-cognate low-affinity binding.

Low-affinity TF-DNA interactions at a non-cognate DNA site could be mediated by various molecular mechanisms: sequence-independent, purely electrostatic TF-DNA interactions; sequence-dependent interactions with low overlap with the consensus motif; or perhaps short-lived stochastic conformational transition of the TF that can affect its sequence-dependent binding affinity. As TFs have to search for their cognate DNA targets among an extensive genomic background, they may encounter many non-cognate sites with sufficient target sequence similarity to allow for binding – either with high or low affinity – thus potentially trapping TF molecules (Gerland *et al*., 2002; Mirny *et al*., 2009). While non-cognate binding of TFs could have a neutral effect on the organism, it could also substantially impair growth for the cell. This could happen either through high-affinity non-cognate binding to a few DNA sites with high similarity to the target sequence, each with a large individual effect; or through low-affinity non-cognate binding to many sites across the genome, each individually with a small effect, but together accruing to a large effect. While large individual effects are more likely to stem from regulatory interference, the mechanistic consequences of widespread low-affinity binding remain largely unknown. The overexpression of one of the *E. coli* nucleoid-associated proteins (NAPs) involved in DNA organization, H-NS, can lead to growth arrest (Spurio *et al*., 1992), suggesting that structural changes in chromosome organization could play a role.

In eukaryotes, which have shorter binding sites, low-affinity binding of TFs seems to be an integral part of gene regulatory function and evolution (Burger *et al*., 2010; Crocker *et al*., 2016; Kribelbauer *et al*., 2019; Shahein *et al*., 2022). In a theoretical study that explored the impact of non-cognate binding we suggested that genomic low-affinity sites impose the existence of a global biophysical constraint, termed “crosstalk” (Friedlander *et al*., 2015). Some forms of TF cooperativity and combinatorial regulation can limit this problem (Friedlander *et al*., 2015), but additional mechanisms, such as out-of-equilibrium proofreading mechanisms and chromatin-based regulation, may also be at work in eukaryotes (Cepeda-Humerez *et al*., 2015; Grah *et al*., 2020; Perkins *et al*., 2025). In prokaryotes, the importance of non-cognate binding to low-affinity sites has been acknowledged in regulatory interference (Sasson *et al*., 2012) but has been rarely investigated, especially with regard to its physiological consequences. Experiments addressing the fitness impact of low-affinity binding are lacking entirely as it is generally difficult to disentangle cognate site-driven effects from non-cognate site-driven effects with native proteins, which rarely have just one cognate binding site (Shimada *et al*., 2018).

## Results

### Experimental setup

We set out to test the consequences of non-cognate TF binding on bacterial growth by expressing TFs lacking known binding sites in native and non-native host cells. The main challenge in addressing this question is that it is not trivial to disentangle physiological effects from non-physiological effects, as many TFs have several cognate binding sites across the bacterial genome and hence multiple or complex biological functions. Based on these considerations, we decided to use phage transcriptional repressors, which are present for long periods at low numbers in bacterial hosts during the lysogenic cycle (Oppenheim *et al*., 2005). As temperate phages likely spend most of their existence as lysogens (Howard-Varona *et al*., 2017), we expect these repressors to be well adapted to the genomic background of their bacterial host. Consequently, these repressors should show few non-cognate binding effects in their hosts — although they may bind cognate binding sites to regulate host genes (Chen *et al*., 2005) — but non-cognate binding could be more prevalent in bacteria that do not serve as host to the particular phage.

Thus, we used TFs from the same protein family from different lambdoid phages, with two being native to *Escherichia coli* (*λ* CI, *434* CI) and one native to *Salmonella enterica* (*P22* C2), and explored their effects on growth in both bacterial hosts. Transcriptional regulation through activation or repression is well-studied for all three TFs (Fattah *et al*., 2000; Ptashne, 2004). The phage repressors bind to their cognate operator sites in the phage genome as a dimer, but can also bind cooperatively to adjacent operators or form short- and long-distance loops (up to 10kbp for λ CI (Priest *et al*., 2014)) involving two to four dimers (Donner *et al*., 1997) (Fig. 1A). This type of TF cooperativity was previously found to facilitate non-cognate binding (Pray *et al*., 1998). Notably for *λ* CI, significant non-cognate binding has been predicted theoretically (Bakk & Metzler, 2004a; Pray *et al*., 1998) and substantiated experimentally (Igler *et al*., 2018), whereas for *P22* C2 binding was lost at few mismatches with the target motif (Igler *et al*., 2018), making it theoretically less likely to bind non-cognate sites (Box 1B).

**Figure 1.**
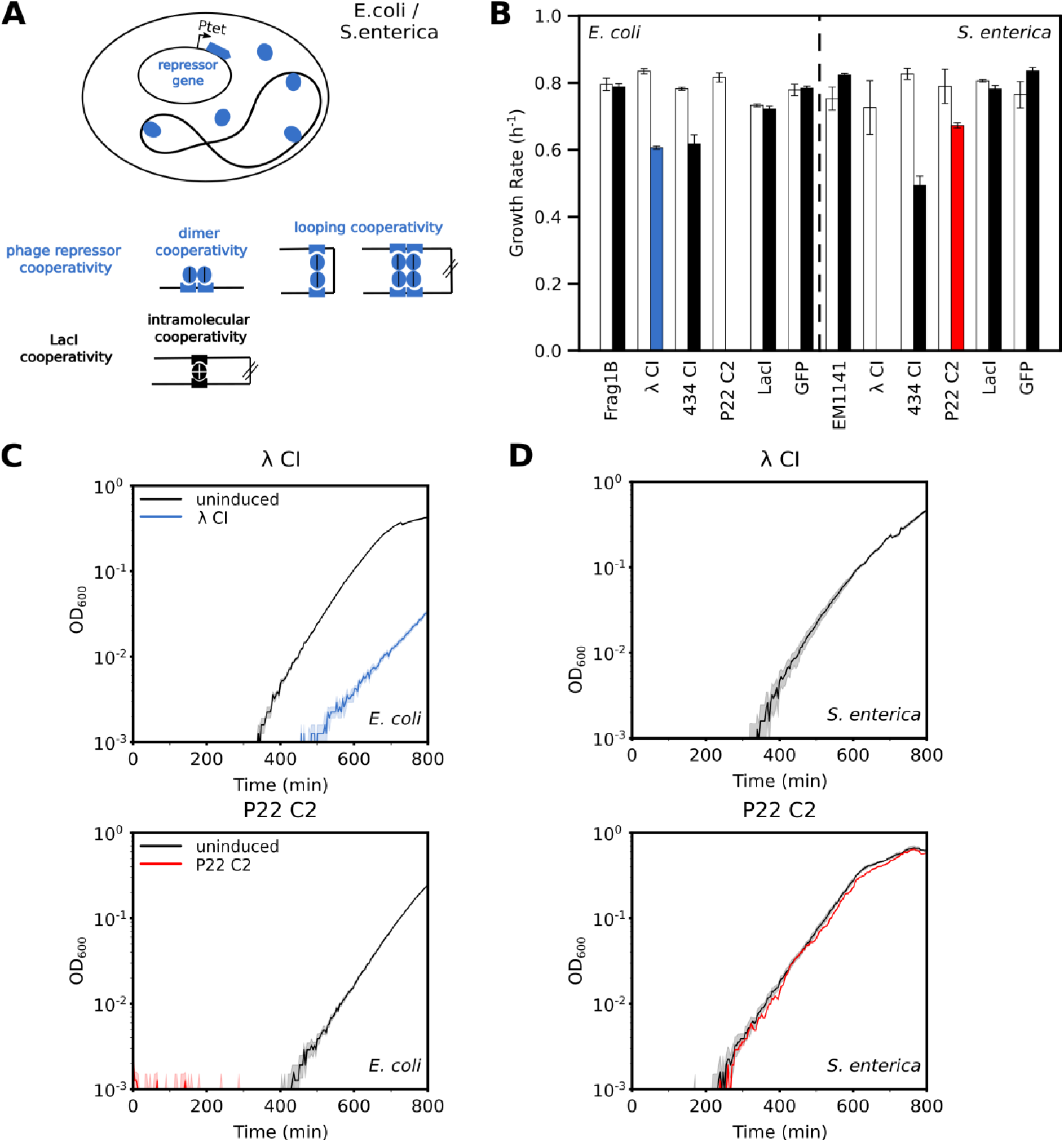
Growth reduction in the presence of repressors in minimal media with glucose. **(A)** The experimental model system with repressors being expressed from a plasmid and their binding cooperativity modes are shown. **(B)** Growth rates per hour (see Methods for calculation) are shown for *E. coli* and *S. enterica* cells in the absence (white) or presence (25ng/mL aTc induction) of λ CI, 434 CI or P22 C2 repressors, LacI, or a GFP reporter (filled bars). **(C,D)** Curves show mean OD_600_ for *E. coli* (C) or *S. enterica* (D) cells in the presence (color) or absence (black) of λ CI (upper, blue) or P22 C2 (lower, red). Note that there is no growth defect for *S. enterica* when expressing λ CI (D, top). Error bars show standard error over 3 replicates (in some cases hidden by the mean).

In addition to the phage repressors, we also chose a well-studied bacterial repressor from a different protein family, LacI, as it is one of the few *E. coli* transcriptional regulators known to have only a single cognate target region at the *lac* operon (consisting of three operator sites) but nowhere else on the chromosome (Shimada *et al*., 2018). LacI uses facilitated diffusion for target search (Elf *et al*., 2007), which suggests that its non-cognate binding tendencies should not significantly affect bacterial growth. LacI is a tetrameric protein and its cooperativity stems from the fact that one repressor tetramer can loop DNA by binding to one DNA site and then the other (Vilar & Saiz, 2005) — as opposed to cooperativity and looping resulting from binding of two separate repressor molecules for the phage repressors (Fig. 1A). The effect of LacI-type cooperativity on non-cognate binding remains unclear.

These four repressors (*λ cI, 434 cI*, *P22 c2* and *lacI*) were each cloned under the control of an inducible promoter (*P_tet_*) on a very low-copy number plasmid (Fig. 1A, S1A). The plasmids were then transformed into *E. coli* and *S. enterica* strains whose genomes do not contain any of the operators for the four repressors (except for one native LacI operator in *E.* coli, Methods). Our *E. coli* strain constitutively expresses TetR (for regulation of *P_tet_*) and LacI, and *S. enterica* constitutively expresses TetR. We tested the effect of constitutive TetR and LacI expression on the growth of our strains and found no effect for either TetR or LacI in *E. coli* (Fig. S1B). We then investigated the impact of non-cognate binding of the four different TFs by measuring growth curves of *E. coli* and *S. enterica* and calculating growth rates as a fitness proxy.

### Growth effects are highly repressor specific

Effects on cellular growth due to the presence of repressors were determined via changes in growth rates (Methods). We observed a wide spectrum of growth behaviors across the four repressors and the two bacterial hosts. For cells grown in minimal M9 media (Methods), the standard media used if not specified otherwise, the induction of *λ* CI resulted in a growth reduction in *E. coli* and halting growth in *S. enterica* cells (Fig. 1B,C,D; S1C; Table S1), whereas the induction of *434* CI repressor reduced growth in both hosts (Fig. 1B). On the other hand, *P22* C2 showed minimal effect in its native host *S. enterica*, while halting growth when expressed in *E. coli* (Fig. 1B,C,D; Table S1). There was no significant impact on growth in both strains for expressing either LacI or a fluorescent protein (Fig. 1B). Thus, three different repressors stemming from the same TF family showed a broad spectrum of growth effects in their native and non-native bacterial hosts. We further explored these growth effects and their causes by focusing on the two well-characterized repressors, *λ* CI and *P22* C2, which are known to have different propensities for binding at DNA sequences far away from their target motif (Igler *et al*., 2018) (Box 1B).

We found that the severe reductions in growth rates translated into a fitness effect in direct competition experiments for both, *λ* CI and *P22* C2. As a ‘neutral’ competitor, we used cells with LacI-expressing plasmids, as LacI was the only repressor showing no significant growth reduction in both bacteria when expressed from the plasmid — even in addition to the LacI that is constitutively expressed in the *E. coli* cells (Fig. 1B, S1B). We added a constitutive YFP-*venus* marker on the LacI-expressing plasmids to determine relative fitness via CFU counts and fluorescence measurements (Fig. S2, Methods). In accordance with the growth rate measurements (Fig. 1,2), repressor expression led to substantial fitness reductions with *λ* CI in *E. coli* and *S. enterica* as well as *P22* C2 in *E. coli* relative to *lacI*+*venus* carriers (Fig. S2, Table S2). As the *venus* marker itself resulted in a slight fitness cost (Fig. S2, Methods), the observed increase in *lacI*+*venus* carrying cells indicates an even more pronounced fitness advantage than measured.

**Figure 2.**
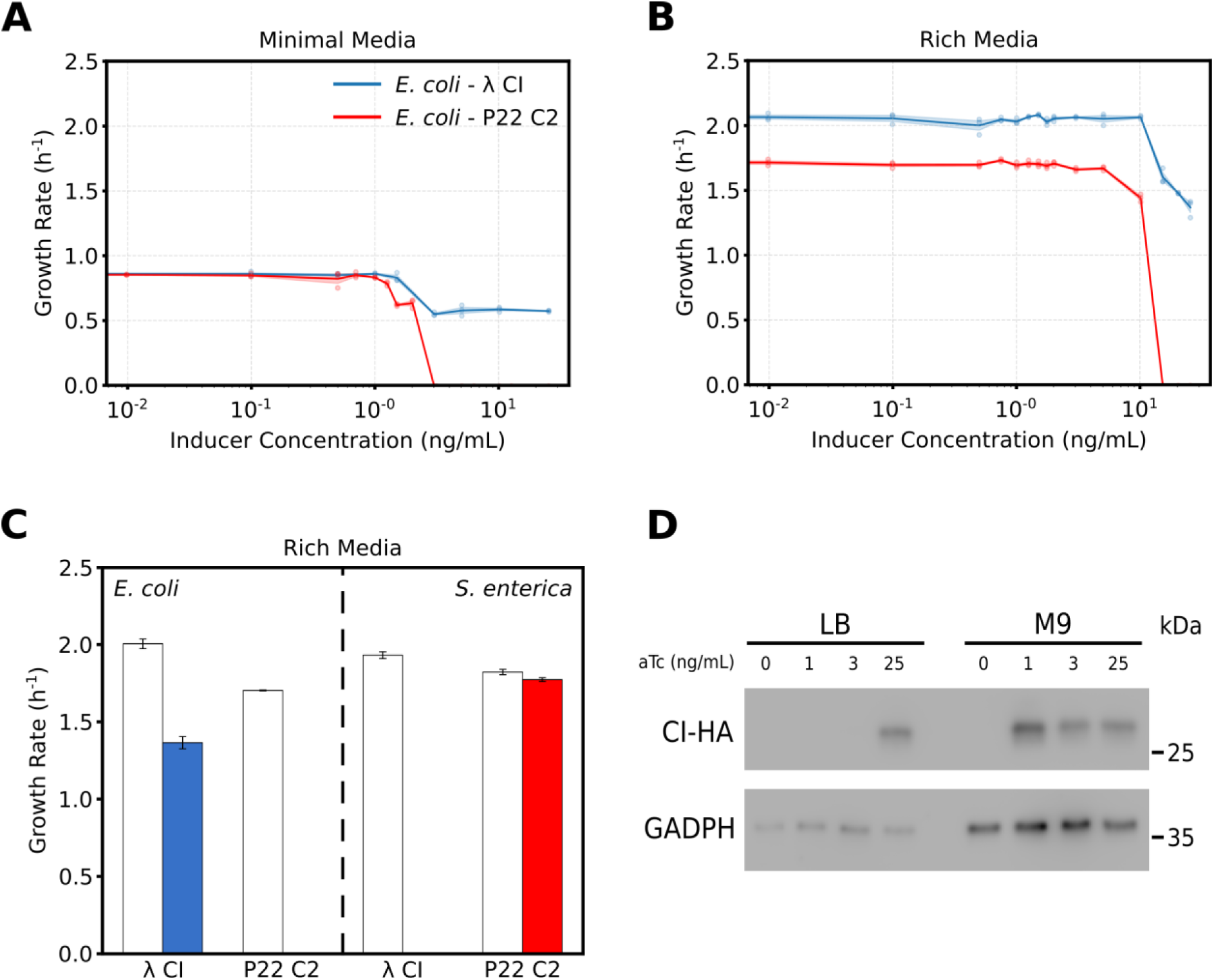
Effect of environment on repressor-dependent growth reduction. (A,B) Growth rates (per hour) are shown for induction of λ CI or P22 C2 in *E. coli* cells over 11 different inducer concentrations (ng/mL aTc) in **(A)** M9 minimal or **(B)** LB rich media conditions. Shaded areas give the standard error over 3 replicates and the solid line is the mean of calculated growth rates. **(C)** Growth rates are shown in the absence (white) or presence (color) of λ CI (blue) or P22 C2 (red) in *E. coli* and *S. enterica* cells in rich media. Error bars show the standard error over 3 replicates. **(D)** Western blot shows how the λ CI-HA protein levels vary in *E. coli* between rich and minimal media for different levels of induction. In LB, λ CI-HA only visibly accumulates under 25ng/mL aTc induction, whereas in M9, 1ng/mL aTc is sufficient to result in accumulation of λ CI-HA. GADPH was used as a loading control. The molecular weight of the proteins is shown in kDa.

### Growth effects are environment and concentration dependent

As the phage repressor levels are likely higher when expressed from our plasmid than in a lysogen, we tested the magnitude of growth reduction at varying repressor induction levels. In *E. coli*, decreasing the induction level of either repressor showed a gradual recovery of normal growth, with *P22* C2 still having an effect at lower induction levels than *λ* CI (Fig. 2A, Table S3). In *S. enterica*, lower induction levels of *λ* CI than in *E. coli* resulted in growth reductions, while *P22* C2 showed no growth changes at any concentration (Table S4).

As a next step, we tested the effect of environmental conditions by growing bacteria carrying *λ* CI or *P22* C2 in LB, a rich medium which allows for faster growth. Growth inhibition in rich media occurs at higher inducer levels than in minimal media (Fig. 2AB), but the qualitative trends remain similar. *P22* C2 shows a growth effect in *E. coli* at lower inducer levels compared to *λ* CI, and no effect in *S. enterica* (Fig. 2BC, Table S4). Interestingly, the growth reduction caused by *λ* CI in rich media occurs at much lower inducer levels in *S. enterica*, but the effect is stronger in *E. coli* (Fig. 2BC; Table S4, S5).

To examine the cause for the dependence of growth reduction on different media, we visualized the presence of *λ* CI tagged with HA via immunoblotting in cells sampled at an OD_600_ ∼0.02 in minimal and rich media with different levels of inducer. In accordance with growth reduction observations, we found that *λ* CI-HA could already be detected at much lower inducer concentrations in M9 than compared to cells grown in LB (Fig. 2D). Lower concentrations of proteins in richer media are expected as faster growth leads to higher dilution of proteins and larger volume of cells (Klumpp, Zhang & Hwa, 2009), which suggests that repressor concentration is correlated with the growth reduction.

### Growth reduction is caused by cooperative, low-affinity binding distributed across the genome

Given the detrimental growth effect of the two repressors in several conditions, we set out to determine its cause. Transcriptional repressors are DNA-binding proteins and could therefore interfere with cellular programs through DNA binding at various non-cognate sites (Chakrabarti *et al*., 2011). To determine the role of TF binding in the observed growth reductions, we used the fact that *λ* CI is one of the best-studied TFs, and thus a range of classical mutants exist that change its binding and conformational properties.

First, we tested the expression of a *λ* CI mutant defective in DNA binding (Nelson & Sauer, 1986) and found that growth rates were somewhat restored in *E. coli*, but greatly restored in *S. enterica* cells (Fig. 3A, Table S1), suggesting that this DNA binding mutant may not be able to bind to the cognate site, but may still able to bind to other DNA, and that the growth defect of *λ* CI in *S. enterica* relies more on site-specific binding. Second, a *λ* CI mutant that cannot form dimers (Whipple *et al*., 1994), which should significantly interfere with the ability of *λ* CI to bind DNA (Weiss *et al*., 1987), completely restored growth in *E. coli* and but only partially restored growth in *S. enterica* cells, suggesting that DNA binding is still required for the growth defect, but additional mechanisms of *λ* CI-induced growth reduction may function differently between the two species. Finally, we tested a *λ* CI mutant defective in cooperativity between repressor dimers – but not in DNA binding itself – which fully restored growth in *E. coli*, and partially restored growth in *S. enterica* cells, suggesting an important contribution from DNA looping or some other form of repressor oligomerization (Fig. 3A, Table S1). This is intriguing as *λ* CI cooperativity and oligomerization are thought to be strengthened by binding to DNA in a sequence-dependent manner (Mazumder *et al*., 2017; Vilar & Saiz, 2005), while the data suggest that cooperativity also appears to play a large role in the absence of cognate sites. Hence, the ability to bind DNA, potentially in a cooperative manner, seems to be central to repressor-mediated growth effects.

**Figure 3.**
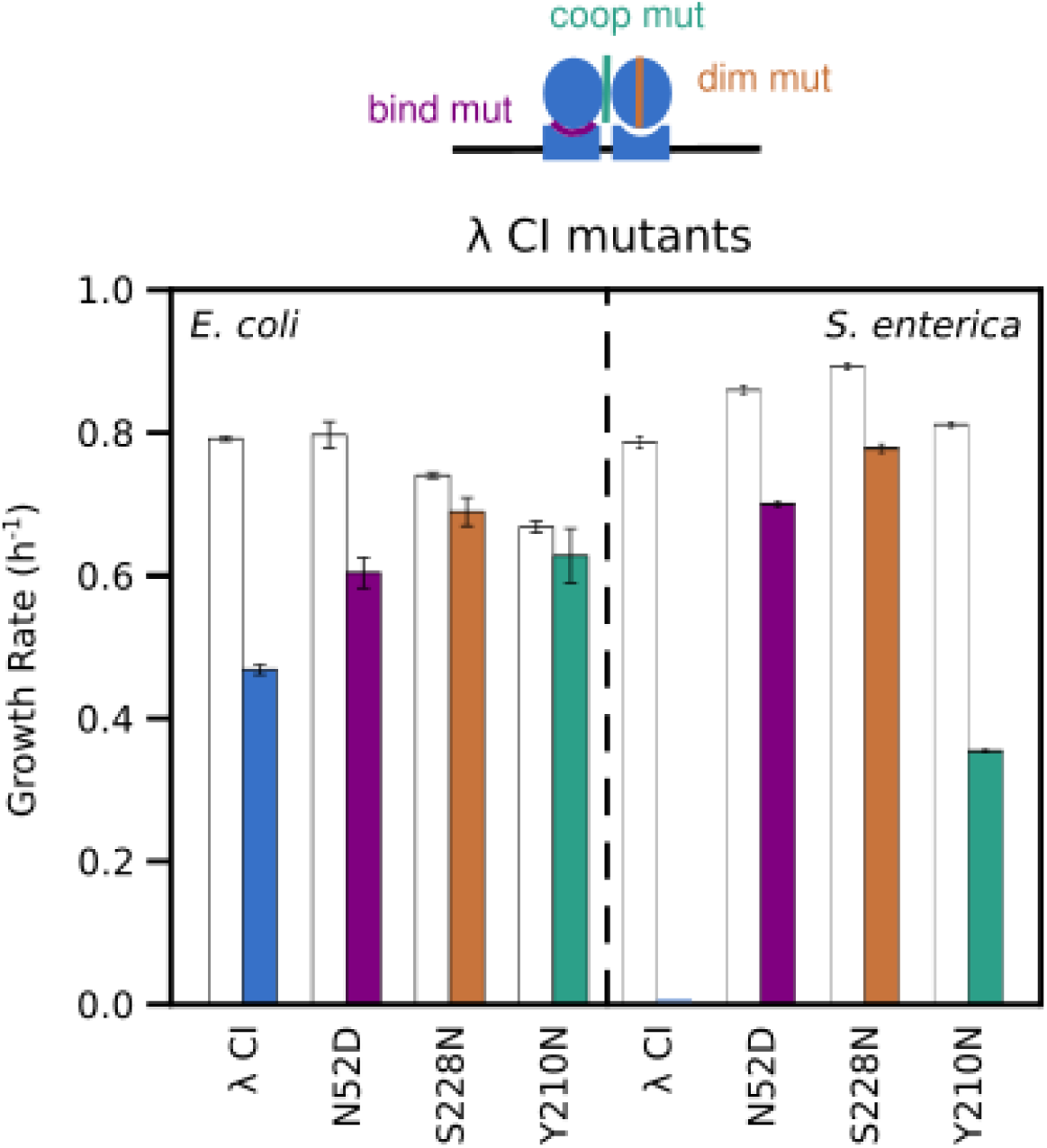
Effect of mutants on repressor-dependent growth reduction. Growth rates (per hour) are shown for *E. coli* and *S. enterica* cells in the absence or presence of a repressor are shown. Comparison between λ CI wildtype (blue), a λ CI binding mutant (purple), a λ CI dimerization mutant (orange) or a λ CI cooperativity mutant (turquoise) in minimal media. Error bars show standard error over 3 replicates.

To determine if the non-cognate binding effects involved either a few essential or many distributed regions of the chromosome, we performed ChIP-sequencing of *λ* CI in *E. coli* and *S. enterica*. As the data did not reveal strong peaks for specific genes in *E. coli*, we investigated overall binding to the genome by averaging data over 1000bp windows. This analysis indicated weak, low-affinity binding at numerous sites all over the *E. coli* chromosome (Fig. 4A, S3A, Table S6) when using a more lenient cutoff than typically used for strong, high-affinity binding. However, we would like to stress that completely sequence-independent binding should not be detectable via ChIP-sequencing, even with our more lenient cutoff (Fig. S4, see Methods section ‘ChIP-sequencing analysis’). In *S. enterica* we found a broad peak (indicating substantial binding in several adjacent genes) and more distributed weak binding regions than in *E. coli*. Interestingly, this broad peak corresponds to prophage regions on the genome that seem to provide binding hotspots for *λ* CI (Fig. 4B, S3B, Table S6). As such binding hotspots were absent in *E. coli* but *λ* CI still showed a growth reduction, we did not consider this finding necessary to qualitatively explain our results, though it may explain the stronger growth reduction in *S. enterica*. None of the peaks for either genome appears to encode an essential gene.

**Figure 4.**
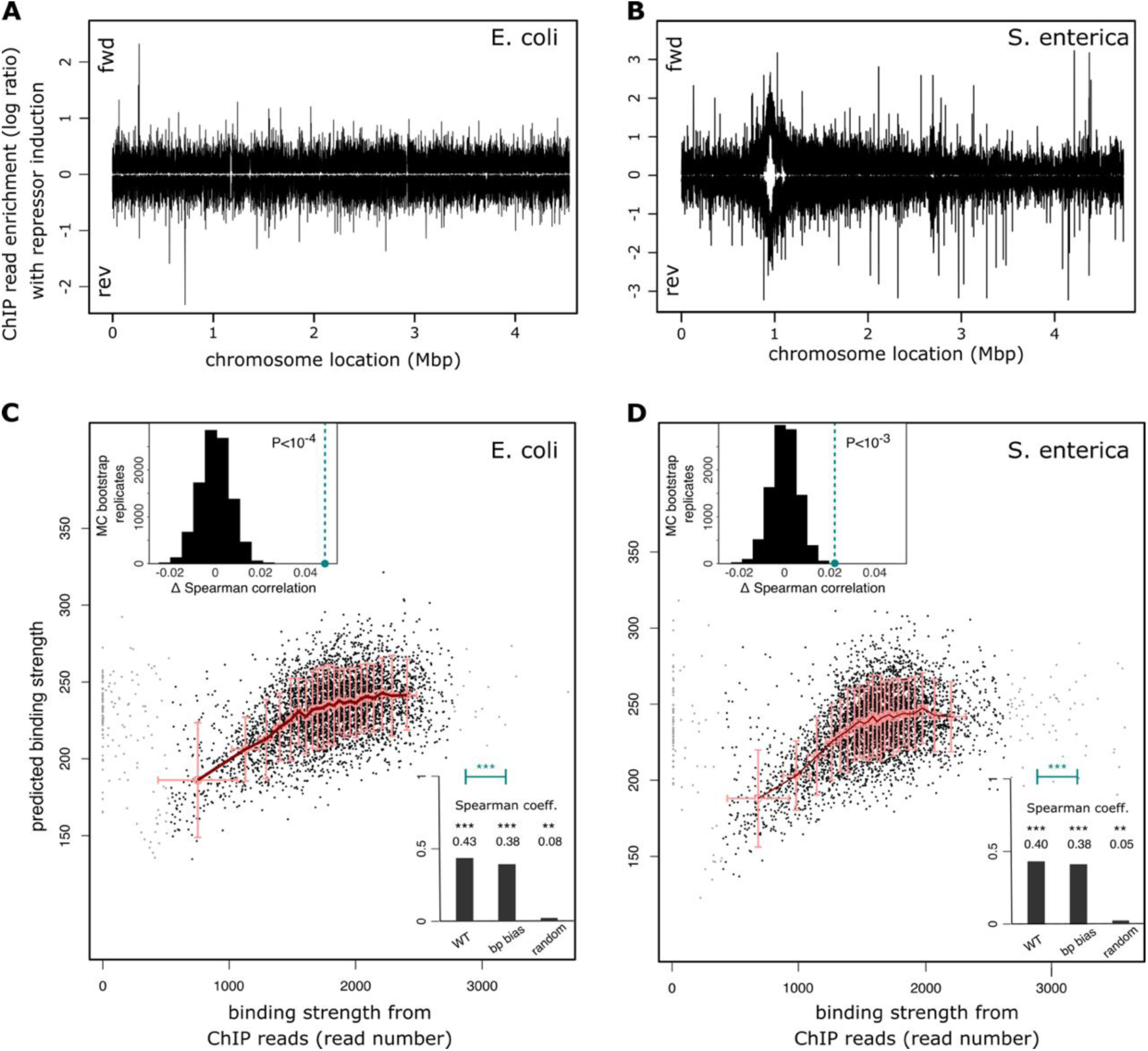
Distributed, low-affinity binding of λ CI across the genomes of *E. coli* and *S. enterica*. (A,B) Distributions of ChIP-sequencing reads for the regions found to be enriched in the experiment with λ CI over the experiment without λ CI are given for the forward and the reverse DNA strand across the **(A)** *E.coli* or **(B)** *S. enterica* genome. Enrichment was calculated within 100bp windows through the log ratio of the scaled coverage (scaled by number of mapped reads) between sample and control ChIP-seq experiments. Number of reads for significantly enriched regions are shown in Fig. S3. **(C,D)** Fit between binding strength predictions of a simple thermodynamic model using the energy matrix for λ CI binding and the ChIP-sequencing reads across 1000bp windows along the **(C)** *E. coli* or **(D)** *S. enterica* genome. In **(D)**, 33 out of 4856 data points showed more than 3600 reads and were omitted for clarity. Lower insets show the calculated Spearman correlations using either the wildtype energy matrix, one that only conserves the λ CI basepair bias (Fits are shown in Fig. S5) or one that has completely reshuffled entries (averaged over 100 permutations). Upper insets in **C** and **D**: Permutation test for the significance of the difference in Spearman correlation between binding predictions using the wildtype energy matrix (first bar in the lower inset) vs prediction with the energy matrix that only conserves λ CI basepair bias (second bar in the lower inset). Black histograms represent the Monte-Carlo-derived null distribution (10^4^ random reassignments of ChIP reads to genomic regions), green dot and line show the true excess Spearman correlation. The correlations for *S. enterica* were not substantially affected by the strong binding peak in prophage regions shown in **(B)**.

As ChIP-sequencing is typically used to determine high-affinity TF binding, there is no established methodology of assessing low-affinity binding in these assays. Here, we used a simple thermodynamic model to predict λ CI binding across the bacterial genome and correlate these model predictions with ChIP-Seq results (see Methods section ‘Thermodynamic model’). The predictions showed a surprisingly high degree of correlation with the number of ChIP-Seq reads (Fig. 4C,D), especially since energy matrix models are believed to perform poorly for low affinity sites (Maerkl & Quake, 2007). Even more surprisingly, most (but not all) of this correlation could be accounted for by an energy matrix that conserved only the λ CI preference for the overall basepair composition (Fig. 4C,D; S5, see Methods section ‘Thermodynamic model’). In contrast, a negative correlation was observed with the overall base-pair preference of *P22* C2 (Fig. S5), which indeed is opposite to that of *λ* CI (Box 1). These results could not be explained by an inherent nucleotide bias of the ChIP-sequencing experiments (Fig. S6), which suggests the correct base-pair composition bias in a genomic sequence could provide *λ* CI with sufficient recognition pattern to bind with low affinity. In agreement with this hypothesis, the GC bias of the stronger *λ* CI cognate operators (*O_R1_*, *O_R2_*, *O_L1_* and *O_L2_*) is 52.94%, which is very close to that of the *S. enterica* genome (52.2%) and somewhat higher than that of the *E. coli* genome (50.8%).

The base-pair composition bias, while important, did not fully account for the agreement between model predictions based on full TF energy matrix and ChIP results. The sequence-dependent contribution to *λ* CI recognition, in excess of the basepair composition bias, was small but remained highly significant in *E. coli* and *S. enterica* (as determined by Monte-Carlo permutation tests; Fig. 4C,D; S7). Overall, our results indicate substantial non-cognate, low-affinity binding due to sequence-dependence and base-pair bias — which is similar to NAP binding patterns (Spurio *et al*., 1992; Yamada *et al*., 1991). Non-cognate binding is facilitated by repressor oligomerization (Pray *et al*., 1998; Sarkar-Banerjee *et al*., 2018), and distributed over the thousands of potential low-affinity *λ* CI binding sites known to be present in the *E. coli* genome (Chakrabarti *et al*., 2011).

### Low-affinity binding leads to cell division arrest

The distributed non-cognate DNA binding demonstrated by the ChIP-sequencing data for *λ* CI is in agreement with the observation that increasing induction of repressor intensifies the growth reduction (Fig. 2A,B). As DNA binding appears to an important factor in growth reduction, the ratio between repressor and potential binding sites, i.e., genomic DNA, may determine the magnitude of the growth effect. If cell doubling time is slower than the time needed for DNA replication, i.e., if the growth rate is less than ∼1.02 doublings/hour in E. coli and ∼1.32 doublings/hour in in S. enterica (Cooper & Ruettinger, 1973), (which is the case in our experiments in minimal media: 0.794(±0.032) doublings/hour and 0.751(±0.0596) doublings/hour respectively), each bacterial cell contains on average only a single chromosome. At faster growth rates, replication cycles are overlapping and daughter cells inherit 2-8 origins at birth, together with partially replicated chromosomes (Nordström & Dasgupta, 2006). Hence, with richer media and consequently faster growth, more DNA will be available to titrate away potentially detrimental non-cognate binding TFs, in addition to the faster growth diluting the concentration of the bacterial repressor (Klumpp, Zhang & Hwa, 2009). ◻This could be an additional factor explaining why much higher induction levels of repressor are required for growth reduction in rich media (Fig. 2): repressors are titrated more due to the presence of more DNA and repressor concentrations are generally lower due to faster dilution as well as larger cell volume.

We set out to test the titration hypothesis by introducing a high-copy number plasmid carrying four cognate *λ* CI binding sites into *E. coli* cells alongside the inducible *λ* CI-plasmid, which should substantially reduce the number of free *λ* CI dimers available for binding. The growth reduction caused by *λ* CI was significantly less in the presence of the high-copy number plasmid at low expression levels (Fig. S8), which suggests that the titration of *λ* CI may alleviate the growth reduction.

The titration hypothesis, together with the Western Blot and ChIP-sequencing results, implies that the overall ratio of chromosomal DNA to repressor proteins could be an important factor in determining the growth effects. This suggests that non-cognate binding may interfere with global cellular functions.

Using microscopy, we found that *λ* CI-expressing *E. coli* cells generally formed long filaments, indicating an inhibition of cell division in these cells (Fig. 5A), which is also characteristic for antibiotics targeting DNA replications and other DNA-related functions (Bollenbach *et al*., 2009). ln the presence of *λ* CI, induction of the stress response promoter *PsulA* is low (Fig. 5B, S9), making the stress response an unlikely cause for the substantial inhibition of cell division (Arends & Weiss, 2004). Furthermore, such a weak stress response will not lead to substantial self-cleavage (i.e., inactivation) of the repressor molecules (Sauer *et al*., 1982) — particularly because *λ* CI becomes a poor substrate for self-cleavage at higher concentrations (Phizicky & Roberts, 1980).

**Figure 5.**
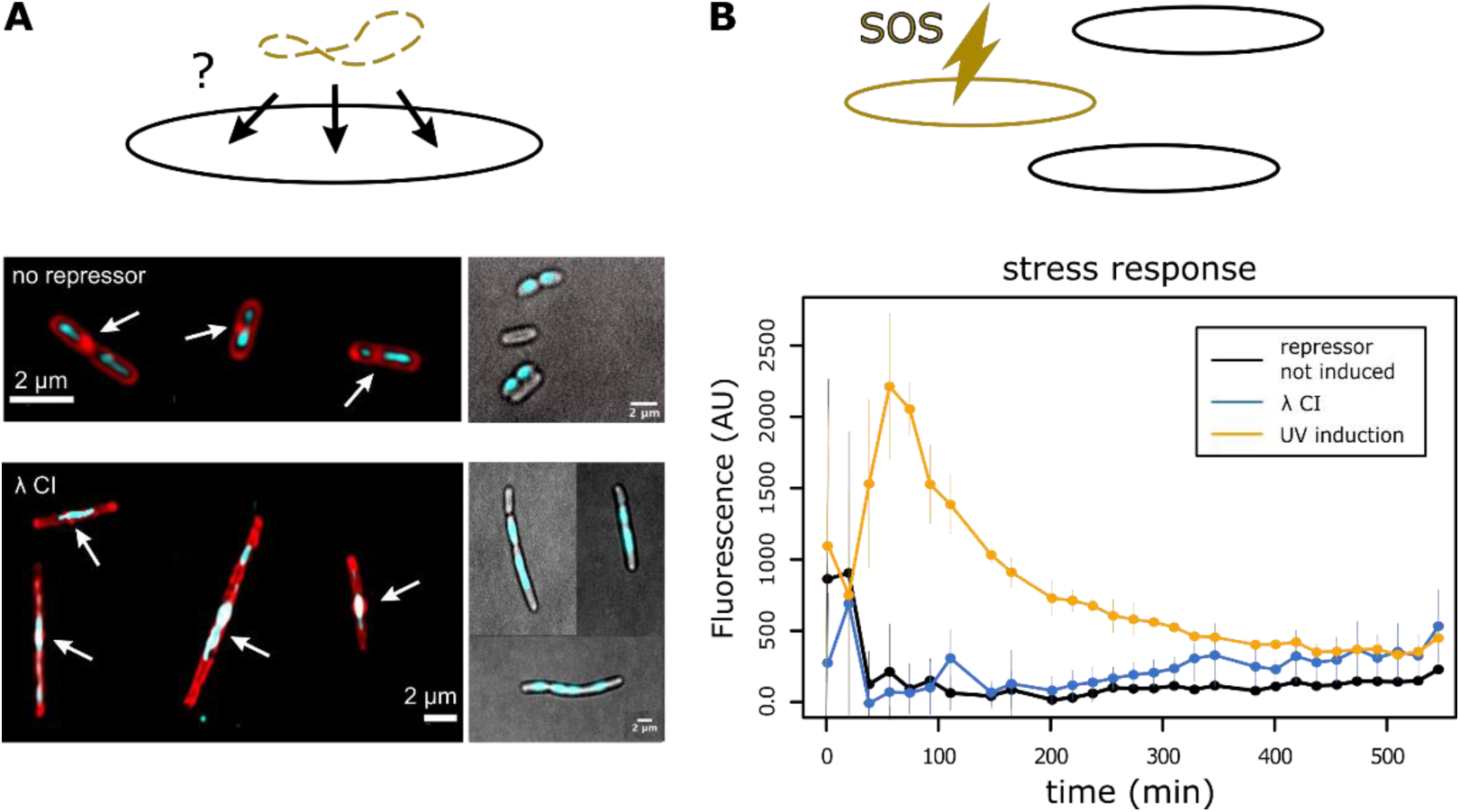
Induction of λ CI interferes with cell division but does not induce the SOS stress response. *E. coli* cells containing the λ CI repressor were **(A)** imaged using dye to visualize the cell membrane in red (Nile Red) and the chromosome in blue (Hoechst) or **(B)** tested for stress response (through the reporter *PsulA*-*yfp*). **(A)** Cells grown in minimal media were imaged using fluorescence microscopy (see Methods) either in liquid media (left) or on an agar pad (right). Whereas chromosomes (blue) are positioned in the middle of the cell in the absence of λ CI (upper), the presence of λ CI (lower) leads to filamentous cells, which often contain several chromosomes, and their positioning overlaps with the middle of the cell including the area of division septa formation (thicker red dots). **(B)** Curves show mean fluorescence for cells that were not induced (black) or cells that were stressed by inducing λ CI expression (blue) or using UV exposure for 30sec (yellow). Error bars show standard error over 3 replicates. (Corresponding mean growth curves are shown in Fig. S9).

Cell division can also be hindered by the presence of genomic DNA at mid-cell (Cambridge *et al*., 2014). Fluorescence microscopy indicates that many *λ* CI-expressing *E. coli* cells that become filamentous contain several chromosomal regions, which are enlarged compared to cells without the repressor; these enlarged regions are often located mid-cell (Fig. 5A). Mid-cell located DNA interferes with the establishment of a division septum, suggesting that it is not FtsZ-ring formation, but a subsequent step in the cell division cascade that is disrupted. Some filamentous cells manage to divide after growing to substantial length as division septa start forming between the chromosomal regions (Fig. 5A). Taken together, non-cognate TF binding could lead to inhibition of cell division by retaining chromosomal DNA at mid-cell.

## Discussion

We investigated the consequences of non-cognate interactions in molecular recognition of DNA by proteins, using three different but related phage TFs and a bacterial repressor. These TFs produced a wide range of effects on host cell growth, from high to no growth reduction; we generally found higher costs in the non-native compared to the native host (Fig.1B, 2C). Taking advantage of the rich and well-established genetics and biochemistry of the classic bacteriophage repressor *λ* CI, we found that its growth reduction likely results from cooperative, low-affinity binding, which interferes with growth by inhibiting cell division. Our data support the hypothesis that binding strategies of prokaryotic TFs are under selection to avoid low-affinity binding to the genomic background (Chakrabarti *et al*., 2011). These results may not generalize to eukaryotes, where weak binding to low-affinity sites can play a functional role (Kribelbauer *et al*., 2019), and the deleterious non-cognate aspects of such binding may be held in check by mechanisms such as silencing or proofreading (Grah *et al*., 2020; Zoller *et al*., 2022, Perkins, Crocker & Tkačik, 2025). These considerations highlight the fundamental differences in gene regulatory design between prokaryotes and eukaryotes, and therefore differing evolutionary constraints (Wunderlich & Mirny, 2008).

For prokaryotes – in contrast to eukaryotes – TF target sites are sufficiently long to allow recognition of single cognate operators with high affinity (Wunderlich & Mirny, 2008). Mismatches to the preferred target sequence lead to progressive loss of binding, but the rate at which this loss accumulates with more mismatches can vary substantially between individual TFs (Igler *et al*., 2018). Among the four transcriptional regulators we analyzed here, *λ* CI combined several properties that together make it a promiscuous DNA binder and thus induce a high fitness cost due to low-affinity binding: strong operator binding (offset) that allows for toleration of more mutations, low mismatch penalties for individual base-pair changes (energy matrix) and strong cooperativity (Box 1B,C) (Igler *et al*., 2018). In contrast, for *P22* C2, the lower offset and higher mismatch penalties make binding to non-cognate sites less likely (Box 1B,C) (Igler *et al*., 2018), producing a significant growth effect only in non-native host cells. The mutant experiments (Fig. 3) argue for a significant contribution of cooperativity – and likely oligomerization – to the growth reduction caused by low-affinity binding, which supports the theoretical finding that TF cooperativity does not alleviate deleterious binding when it stabilizes cognate *as well as* non-cognate binding (Friedlander *et al*., 2015). LacI’s lack of intermolecular cooperativity makes it less likely to bind non-cognate sites, suggesting an adaptation for high affinity to only few cognate sites - as opposed to most *E. coli* regulators (Shimada *et al*., 2018). However, it is important to note that cooperativity’s contribution to growth reduction may be due to other mechanisms. The continuum of growth effects we observed for only four different TFs reveals that the complex combination of binding cooperativity, offset, and TF concentration determines the propensity for non-cognate binding of a TF and must be tuned to avoid fitness costs. Therefore, considering binding to low-affinity sites may be crucial in choosing TFs for synthetic biological systems to avoid global toxicity effects, as well as unwanted TF titration, which can affect target gene regulation (Brewster *et al*., 2014; Sasson *et al*., 2012).

The magnitude of the growth reduction depends on a repressor’s ability to bind non-cognate low-affinity DNA sites, as well as on a repressor’s relative ratio to the total amount of DNA within the cell. Slow cell growth compounds the effect as cells contain less DNA that would provide potential sites for low affinity binding and repressor titration, as well as having a higher repressor concentration than at fast growth (Klumpp, Zhang & Hwa, 2009). Hence, the slower growth rate in minimal media could allow for greater accumulation of repressor, which in turn slows growth, resulting in negative feedback between repressor accumulation and growth rate, which would explain the difference in repressor concentration between minimal and rich media. This phenomenon is reminiscent of growth bistability in drug resistant bacterial cells, which is caused by feedback between the growth rate and the speed of counteracting toxic agent (Greulich *et al*., 2015). In our system, the repressors can be seen as “toxic agents”, which are “counter-acted” by dilution or titration if cells manage to start growing fast enough; otherwise, they are growth-arrested.

Additionally, stress tolerance could be higher under optimal growth conditions as found in rich media (Pleška *et al*., 2016). However, it does not seem likely that media-specific genes are targeted, as ChIP-sequencing generally revealed distributed, low-affinity binding all over the chromosome (Fig. 4). Rather, inhibition of cell division seems to result at least partially from nucleoid localization at mid-cell (Fig. 5). Clearance of the division site is impeded if sister chromosomes fail to segregate (Cambridge *et al*., 2014; de Boer, 2010), which may be caused by the formation of “bridges” between cooperatively bound repressors, holding the sister chromosomes together. As found for *λ* CI, even non-cognate binding can increase and stabilize DNA loops through the formation of higher-order oligomers (Manzo *et al*., 2012; Sarkar-Banerjee *et al*., 2018). This provides an additional explanation on why repressor cooperativity is necessary to observe growth effects: in addition to stabilizing non-cognate binding, cooperativity is required to form the “bridges” that inhibit chromosome segregation and thus inhibit cell division. In such a case, the relative ratio of repressor to DNA would govern how tightly the chromosomes are bound together, and sufficiently low repressor to DNA ratios would be unable to prevent the chromosomes from segregating.

Generally, the compaction of nucleoids seems to be a crucial factor for preventing the completion of division septa through nucleoid occlusion (Sun & Margolin, 2004). Intriguingly, a very similar growth effect has been found with H-NS, one of the NAPs responsible for chromosome organization and compaction in *E. coli* (Spurio *et al*., 1992; Yamada *et al*., 1991): H-NS overproduction drastically reduced cell viability, which seemed to be related to the formation of higher-order H-NS oligomers (Spurio *et al*., 1997) – a state that is favoring its ability to form bridges between DNA regions (Verma *et al*., 2019). As cell growth, shape, division and DNA replication are thought to be tightly linked in complex and poorly understood ways (Amir *et al*., 2019), a mechanistic explanation of the observed division inhibition is presently not possible, but prior studies on H-NS combined with our findings make the case for the existence of general constraints on DNA-binding proteins.

Our results suggest that the inherent ability of DNA-binding proteins to occupy non-cognate DNA regions can pose, in addition to potential regulatory interference (Sasson *et al*., 2012), a substantial challenge for host cell fitness overall – particularly considering facilitating conditions like cellular crowding (Zabet & Adryan, 2013), horizontal gene transfer (Baltrus, 2013) and mutations that alter the binding affinity of a protein. This challenge stems from the fundamental limits to molecular recognition that are set by the biophysics of molecular interactions (Friedlander *et al*., 2015) and could lead to various non-cognate effects on the physiology of the cell. The four transcription factors used here show a variety of non-cognate effects, which could indicate different selection pressures that have been acting on their binding and cooperativity characteristics as well as different potential for being tolerated in a particular environment. There might also be a variety of mechanisms underlying the growth phenotype, as suggested by the ChIP-sequencing results for *λ* CI: the additional strong peak region found in *S. enterica* could indicate an additional effect of higher-affinity sites and may explain the stronger growth reduction in minimal media as compared to *E. coli* (especially at low repressor concentrations). A potential for high fitness costs, even in native environments as seen here with *λ* CI and *434* CI, can limit the number and binding affinity of promiscuous DNA-binding proteins in the cell (Friedlander *et al*., 2015). Less promiscuous binders such as *P22* C2 are only detrimental in non-native environments. Hence, the influx of foreign genes through horizontal gene transfer could be considerably impaired through non-cognate binding effects, as horizontal gene transfer and non-cognate binding are likely to occur under slow growth conditions, where already low levels of transcription factors might be costly (Fig. 2, S2). As the phage repressors we used originate from temperate phages, interference with host cell growth can limit their potential host range regarding successful establishment of lysogeny. Considering that phage repressor concentrations are kept low during lysogenic cycles, the selection pressure to reduce low-affinity binding might generally be weak, which would explain the diversity in growth effects we observed with related phage repressors. Phage TFs expressed during lysogeny have however been suggested to be able to change bacterial cell physiology (Derdouri *et al*., 2023). More generally, experimentally uncovering the fundamental biophysical constraints imposed by the non-cognate interactions of TFs is difficult, as TFs with many cognate targets need to recognize a diversity of sequences and by default affect many cellular functions, while single target TFs are rare (Shimada *et al*., 2018). This is what ultimately motivated our choice of focusing our experiments on phage repressors and LacI.

We have shown experimentally that non-cognate recognition of biomolecules can constitute a limiting factor for cellular function and evolution due to the fundamental biophysical constraints on protein-DNA interactions. However, these costs could be counter-balanced by increased TF robustness to target site mutations or higher evolvability, precisely because interactions can be more easily formed at low affinity (Friedlander *et al*., 2017; Igler *et al*., 2018; Shepherd *et al*., 2023; Taylor *et al*., 2022). For example, a TF could co-opt regulation of a non-cognate gene – even if only to a small degree – that provides an advantage in a certain environment, which can subsequently be refined by evolution, a process which has been observed experimentally (Taylor *et al.,* 2015). This opens up wider questions about the interplay of costs and benefits of low-affinity molecular interactions, especially when these interactions also serve as drivers of evolution (Friedlander *et al*., 2017).

## METHODS

### Strains

*E. coli* Frag1B is an MG1655-derived strain that constitutively expresses TetR (P_N25_ promoter) and LacI (P_laci_^q^ promoter) together with a spectinomycin resistance from a cassette that was inserted twice into the chromosome at the λ attachment site (Le *et al*., 2005).

*S. enterica* EM1141 is a derivative of *S. enterica* LT2 (a gift from the M. Erhardt lab). It does not carry any lac repressor and has a tetracycline resistance cassette (tetra) inserted near the P22 attachment site.

### Plasmids

#### Wild-type repressors

The phage repressors *λ* CI, *P22* C2, *434* CI or the bacterial repressor LacI were cloned under the control of a P_LtetO-1_ promoter in a low copy number *kan^R^* plasmid (pZS21) by using the restriction enzymes KpnI and XbaI (Lutz & Bujard, 1997). In control experiments the phage repressor on the plasmid was replaced by a fluorescence marker gene (*gfp*). The low copy number plasmids used are under stringent replication control, which is linked to chromosome replication (Kües & Stahl, 1989).

#### Sponge plasmid

Titration of λ CI was tested by transforming *E. coli* cells containing the pZS21-*λ cI* plasmid with the sponge plasmid: a compatible, high-copy number chloramphenicol resistant plasmid (pZE3, 50-70 copies, Lutz & Bujard, 1997) with the natural λ CI operators, *O_L1_* (x2), *O_L2_* (x2), cloned between the XhoI and HindIII sites. The control plasmid is identical, but with all four the operator regions replaced with random DNA. Four operators per plasmid and 50-70 plasmids per cell results in approximately 200-280 operators per cell. The sponge plasmid insert is TCGAGTATCACCGCCAGTGGTATTTATGTCAACACCGCCAGAGATATTTATGTTTTATGTTA TCACCGCCAGTGGTATTTATGTCAACACCGCCAGAGATA, and the control plasmid insert is TCGAGGCGCGAAAGTTCGTACCTTTATGTGCTCATTCACTTGGTTCTTTATGTTTTATGTCG AAGCCTGTCCTGATATTTATGTTATGAATCCAAACGATA.

#### *λ* CI-HA

A haemagglutinin tag (TATCCGTATGATGTTCCGGATTATGCATGA, YPYDVPDYA, Field *et al*., 1988) was cloned on the carboxy-terminal end of the the *λ* CI gene in the pZS21- *λ* CI plasmid.

#### *λ* CI mutants

Based on previous studies, we cloned three different λ repressor mutants into the same low copy number plasmid (pZS) under the control of a *P_LtetO-1_* promoter: a repressor mutant that is defective in its ability to bind DNA (N52D) (Nelson & Sauer, 1986), a repressor mutant that cannot form dimers (S228N) (Whipple *et al*., 1994) and hence can no longer effectively bind DNA, and a mutant that can dimerize but not form higher-order oligomers, i.e. that cannot bind cooperatively (Y210N) (Whipple *et al*., 1994). Mutations were introduced using site- directed mutagenesis. The function and stability of the proteins has been shown previously (Nelson & Sauer, 1986; Whipple *et al*., 1994), as well as that their production levels are not different from wildtype repressor (Whipple *et al*., 1994).

### Growth measurements

All cultures were grown overnight at 37°C in M9 medium supplemented with 1mM thiamine, 0.4% glucose and 25μg/mL kanamycin (except when specified differently), then diluted 1:1000 in the same media and grown to OD_600_ 0.1. Cultures were then used to dilute (1:1000) 3 replicates without inducer and 3 replicates with different levels of aTc in 96 well plates and were grown and measured at 37°C under shaking at 220 rpm in a Bioscreen Spectrophotometer. Populations were measured (OD_600_) every 4 minutes for up to 24 hours. Population growth in the spectrophotometer was compared to population growth in a culture tube shaken in an incubator at 37°C with sampling via plating for CFU counts after 0, 300, 360, 420 and 480 min (2 biological and 3-6 technical replicates per time point).

Growth windows for growth rate calculation were chosen to be within an OD_600_ range of 0.018 and 0.135. Growth rates were calculated for each replicate individually using linear regression.

To monitor induction of the stress response, we used a strain from Pleška et al. (2016) with a fast-maturing yellow fluorescent protein (YFP, (Nagai et al., 2002)) fused to the promoter of sulA (PsulA-yfp), which was placed on the chromosome using lambda red recombineering (Costantino & Court, 2003). sulA is strongly upregulated as a part of the stress response (Friedberg et al., 2005). The PsulA-yfp strain was then transformed with the pZS21-λ cI plasmid. We checked induction of the reporter by exposing cells to UV light for 30 seconds using the UV transilluminator function of the BioRad GelDoc XR+ station. These cells were diluted 1:1000 from overnight cultures in M9 medium supplemented with 1mM thiamine, 0.4% glucose and 25μg/mL kanamycin, grown for 2h before UV exposure, and then grown and measured in the plate reader after exposure.

### Competition assays

In order to distinguish between strains carrying a phage repressor (showing growth reduction) or the bacterial repressor LacI (not showing growth reduction), we used the pZS plasmid carrying *lacI* (under the control of P_LtetO-1_), together with a constitutively expressed fluorescence marker (*venus*) cloned in the opposite direction upstream of P_LtetO-1_. Venus was expressed from a mutated version of P_R_, which abolishes *λ* CI binding affinity in the *O_R1_* operator (Flashman, 1978; Sarai & Takeda, 1989) (the mutated *O_R1_* sequence used is: TGCCTTAATACTGGATA) – and does not contain *O_R2_* or *O_R3_* - and is therefore constitutive (‘P_constitutive_’). The fluorescence marker was placed on the LacI-carrying plasmid as LacI showed no growth effect in *E. coli* or *S. enterica* (Fig. 1B, Table S1). For E. coli that means that there was no additional effect of expressing more LacI than from the chromosome. As chromosome expression of LacI is constitutive, induction of TF expression from plasmids still occurs under the same genomic background conditions and can be compared in direct competition. As there was no growth effect of LacI-carrying plasmids, bacterial cells did not show filamentation and hence there was no effect of cell morphology changes on fluorescence. There was a slight fitness cost due to the presence of the fluorescence marker on the plasmid in the presence of 0.8ng/mL aTc induction (Fig. S2), which however only strengthens our findings that LacI-expressing cells increase in competition with phage repressor-expressing cells despite this cost.

Overnight cultures of clonal *E. coli* cells carrying pZS21-*λ cI* or carrying pZS21-*lacI* were adjusted to ∼10^8^ cells and mixed 1:1 with a culture of the same cell density carrying pZS21- *lacI*-P_constitutive_*-venus.* Cultures were grown in M9 minimal media supplied with 0.4% glucose, 1mM thiamine, 25 μg/mL kanamycin and 0.8ng/mL aTc at 37°C in a shaking incubator and sampled via serial dilution plating after 0, 300, 360, 420 and 480 minutes. Cells were distinguished based on the fluorescence marker. For the *λ* CI versus LacI+Venus competition, we used 3 biological replicates and 3-6 technical replicates each. For the control LacI versus LacI+Venus competition, we used 4 technical replicates each for one induced and one uninduced experiment. Selection coefficients w were calculated using w=ln[(B_t_/B_0_)/(V_t_/V_0_)], where B_t_ represents CFU counts of cells with *λ* CI or LacI and V_t_ represents cells carrying LacI+Venus at t=480min.

We further used fluorescence measurements as a proxy for abundance to compete a larger number of host strain (*E. coli* or *S. enterica*) – plasmid (pZS21-*lacI,* pZS21-*λ cI*, pZS21-*P22 c2*, pZS21-*lacI-*P_constitutive_*-venus*) combinations. Strains containing an unmarked repressor plasmid were mixed 1:1 with a strain carrying pZS21-*lacI*-P_constitutive_*-venus*, diluted 1:100 into fresh medium and grown in 96-wellplates for 10h. Fluorescence was measured every 30min and compared between cultures that were induced with 25ng/mL aTc and cultures that were not induced.

### Immunoblotting

Overnight cultures of Frag1B::pZS21-*λ cI-HA* grown at 37°C under shaking at 220 rpm in M9 supplemented with 0.4% glucose, 1mM thiamine and 25 μg/mL kanamycin or LB supplemented with 25 μg/mL kanamycin were diluted in 50mL of fresh media to an OD of 0.005. Samples were induced with 0, 1, 3, or 25 ng/mL aTc and grown at 37°C under shaking at 220 rpm to an OD of 0.02-0.03. 40mL of culture was pelleted at 4°C at 8000g for 10 minutes, washed in 40 mL of PBS, repelleted under the same conditions, and resuspended in 1mL PBS. Samples were then pelleted at 4°C at 21130g for 10 minutes, resuspended in 200μL PBS and stored at -70°C. OD measurements were done using a Hitachi U-5100 Spectrophotometer. Samples were normalized to an original OD of 0.02 with PBS, and mixed in a 3:1 ratio with 4x Laemmli sample buffer and incubated at 95°C for 20 minutes. 5μL of each sample was separated by electrophoresis on a 12% Mini-PROTEAN® TGX™ Precast Protein Gel (Bio-Rad, 4561046), and transferred to a polyvinylidene fluoride (PVDF) membrane using the Trans-Blot Turbo Transfer System (Biorad). The transferred PVDF membrane was blocked at room temperature for one hour with gentle agitation in Tris-buffered saline containing 0.1% Tween 20 (TBST) and 1.5% skim milk, then stained with HA Tag Monoclonal Antibody (2-2.2.14) (Invitrogen, 26183-HRP) at 4°C with gentle agitation overnight. The membrane was washed thrice for 10 minutes in TBST, treated with Supersignal™ West Fempto Maximum Sensitivity Substrate (Thermo Scientific, 34096), and visualized with an Amersham™ Imager 600 RGB (GE) with an exposure time of 30 sec. To visualize the loading controls, the membrane was stripped by incubating with Abcam mild stripping buffer (15 g/l glycine, 1 g/l SDS, 1% Tween 20, pH 2.2) twice for ten minutes at room temperature, washed with PBS twice for 10 minutes followed by washing with TBS twice for 10 minutes, reblocked at room temperature with gentle agitation in TBST + 1.5% skim milk, and restained with GAPDH Loading Control Monoclonal Antibody (GA1R) (Invitrogen, MA5-15738-HRP) for 2 hours at room temperature with shaking. The membrane was treated with Supersignal™ West Fempto Maximum Sensitivity Substrate (Thermo Scientific, 34096) and visualized with an Amersham™ Imager 600 RGB (GE) with 6 sec exposure.

### ChIP-sequencing analysis

To perform ChIP-sequencing experiments, we used the HA-tagged *λ* CI plasmid. HA-tagged *λ* CI showed the same growth phenotype as wild-type in minimal and rich media (Fig. S10, Table S7). Samples from strains grown in the presence or absence of *λ* CI were prepared according to (Waldminghaus & Skarstad, 2010).

For ChIP-seq, cells were grown at 37°C to an OD600 of about 0.1-0.2 in 50 mL LB before 27 μl of formaldehyde (37%) per mL medium were added (final concentration 1%). Crosslinking was performed at slow shaking (100 rpm) at room temperature for 20 min followed by quenching with 10 mL of 2.5 M glycine (final concentration 0.5 M). Cells were collected by centrifugation and washed twice with cold TBS (pH7.5). After resuspension in 1 mL lysis buffer (10 mM Tris (pH 8.0), 20% sucrose, 50 mM NaCl, 10 mM EDTA, 10 mg/mL lysozyme) and incubation at 37°C for 30 min followed by addition of 4 mL IP buffer, cells were sonicated at 4°C in 1.5 mL tubes with 48 cycles (30 sec. sonication, 30 sec. cooling) with Bioruptor Plus from Diagenode. After centrifugation for 10 min at 9000 g, 800 μl aliquots of the supernatant were stored at -20°C or processed directly. 800 μl of sonicated cell extract was incubated with 20 μl protein A/G agarose beads (Ultralink) and the different antibodies at 4°C overnight with slow mixing. Agarose beads were collected at the bottom of a 1.5 mL Eppendorf tube by centrifugation. The supernatant was then removed by pipetting. Washing was done by adding 700 μl buffer to the beads and rotation at room temperature for three minutes with subsequent collection of the beads by centrifugation as above. Washing was performed with the following buffers (IP buffer two times all others one time): IP buffer (50 mM HEPES-KOH pH 7.5, 150 mM NaCl, 1 mM EDTA, 1% Triton × 100, 0.1% Sodium deoxycholate, 0.1% SDS), IP buffer with 500 mM NaCl, wash buffer (10 mM Tris pH 8.0, 250 mM LiCl, 1 mM EDTA, 0.5% Nonidet-P40, 0.5% Sodium deoxycholate) and TE. For elution, 100 μl elution buffer (50 mM Tris (pH 7.5), 10 mM EDTA, 1% SDS) was added to the column with the beads, incubated in a 65°C water bath for 10 min and centrifuged as above. DNAs were incubated with RNase A (50 μg/mL) for at least 90 min at 42°C. To reverse the cross link 80 μl TE and 20 μl proteinase K (20 mg/mL) were added and samples incubated for 2 h at 42 and 6 h at 65°C. DNA was purified with phenol/chloroform. The control DNA was taken from the supernatant resulting from centrifugation of the precipitated chromatin beads and processed further as the immunoprecipitated DNA after elution.

Library preparation and Illumina Sequencing was performed at the VBCF NGS Unit (www.vbcf.ac.at). The obtained sequencing data was analyzed using Galaxy and RStudio. Reads were aligned against a reference genome (E. Coli: NC_000913.2, Salmonella: AE006468.2) using bowtie (v1.1.2) and --wrapper basic-0 -q -p 6 -S -n 2 -e 70 -l 28 --maxbts 125 -k 1 -m 1 parameters. Alignment statistics are given in Table S6.

We performed peak calling by examining partially overlapping (50% overlap) sliding windows of 100bp each. Within each 100bp window, we calculated the log ratio of the scaled coverage (scaled by number of mapped reads) between sample and control ChIP-seq experiments while permitting a maximum of five mapped reads to share the same genomic coordinates. Enrichment, i.e. positive log ratios, are plotted in Fig. 4 (A,B). We calculated strand-wise p-values for enrichment by first resampling scaled read coverage within each fragment and then randomly partitioning them to calculate enrichments. Finally, we identified bound regions to be those with positive enrichment scores on both strands with a Benjamini-Hochberg false-discovery rate of less than 30% as we were looking for binding with low affinity and ChIP-binding data was previously found to be highly informative for a wide range of affinity profiles (Tanay, 2006). We estimated the false-discovery rates to assess significance of every peak. To account for local biases within the genome (Zhang *et al*., 2008), we utilized non-overlapping 2kb genomic fragments to estimate FDRs. Different choices of sliding windows (25bp, 50bp, 100bp), maximum number of reads at a specific genomic coordinate (3, 5, 10) as well as genomic fragment size (1000bp, 2000bp, 5000bp) identified similar sets of peaks. In Fig. S3, we plotted the read-depth across the genome for the regions that showed significant enrichment in this analysis and for comparison we plotted the read-depth for not significantly enriched regions. As control for our ChIP-sequencing procedure and analysis we used antibodies against SeqA, which gave the expected peaks as published previously (Waldminghaus & Skarstad, 2010).

We calculated the nucleotide composition of the sequences underlying enriched regions in ChIP-seq data for both bacterial species (Fig. S6). To test for sequence composition bias in these enriched regions, we sought to test if the sequence compositions of the enriched regions were significantly different compared to the rest of the genome. To this end, we randomly selected 50 genomic regions with at least 5kbp distance between them. We then calculated the nucleotide composition of these randomly selected regions and by repeating this procedure 1000 times, generated a null distribution for sequence composition of randomly selected genomic regions. Similarly, we calculated the di-nucleotide composition (with 1 bp overlap) of the same randomly selected genomic regions and compared it to that of the enriched regions.

### Thermodynamic model

The unfiltered number of reads within 1000bp windows was compared with the predicted binding by calculating binding energy at each genome position (using a sliding window approach) from the λ CI offset (i.e. the energy difference between the repressor being bound with high affinity to a cognate operator and being free in solution (Koblan & Ackers, 1992)) and the energy penalty as given by the λ CI energy matrix (Sarai & Takeda, 1989). Lower energies result in stronger binding, meaning positive energy penalties decrease binding affinity (note that negative penalties could increase binding over the one seen with λ CI wildtype operator sites). Binding strength was calculated using 1/(1+exp(E-µ)), with E being the calculated binding energy, as described above and used in (Igler *et al*., 2018), and µ being the chemical potential, which we optimized to give the highest Spearman correlation fit (2.6 in *E. coli* and 2 in *S. enterica*). For comparison with the number of ChIP-sequencing reads, calculated binding strength was summed over the same genomic 1000bp regions (results shown are considering binding only to the forward strand, for ChIP reads and predictions, but the results remain the same when using both strands). In Fig. 4 (C,D) we plot a non-parametric, non-linear relationship estimate between the predicted binding energy and the ChIP-sequencing reads obtained from a series of conditional medians. To investigate the dependence of the correlation between the affinity predictions and the ChIP-sequencing reads on the structural versus the sequence information contained in the energy matrix, we repeated the analysis with i) a matrix of the same size that conserves only the ACGT bias of the λ CI energy matrix (each row contains the average value of that row) or ii) matrices that had completely reshuffled entries. For the latter the average correlation was taken over 100 permutations.

To assess the importance of sequence motif information versus nucleotide (GC) bias, we used a Monte-Carlo permutation test: We calculated the difference between Spearman correlations of ChIP reads with binding prediction using the wildtype energy matrix vs binding prediction using the energy matrix that only conserves λ CI basepair bias, for the true ChIP read assignment, and 10^4^ random read assignments (null distribution). We found an overall strongly significant difference in *E. coli* and lower significance in *S. enterica* (Fig. 4C,D), even though the effect size was small. This means that while most of the measured ChIP signal can be accounted for by a TF model that predicts binding based on the nucleotide content of genomic fragments alone, there is a small but highly significant residual ChIP binding signal that requires the full binding site preference (energy matrix), not just single nucleotide bias, to be explained. Furthermore, we examined the influence of GC content by repeating the Monte-Carlo permutation test for genomic sequences of a specific GC %. Here, we found only a significant motif contribution for the 49% bin in *E. coli* (Fig. S7).

Additionally we used the offset and energy matrix for LacI (Barnes *et al*., 2019; Vilar & Saiz, 2005) and P22 C2 (Hilchey *et al*., 1997) to predict binding and calculate the Spearman correlation with the λ CI ChIP-sequencing reads (Fig. S5). The magnitudes of the Spearman correlations for LacI and P22 C2 predictions are similar to the ones with the λ CI basepair bias, but negative (inverse) for P22 C2. Basepair bias of the energy matrices was calculated as the sum of the average A and T preference minus the sum of the average G and C preference.

### Microscope fluorescence measurements

Imaging of cell membranes and DNA positioning was done using a TIRF Leica DMI6000B (inverted) microscope with 1) an Andor iXon EM CCD camera (front illuminated, 8x8 square micron pixel size) and a 100x 1,47Na Oil HCX Plan Apo objective, giving an effective pixel size of 64nm/pixel; and 2) a Nikon Ti-2 microscope with a 100× oil immersion objective (Plan Apo λ, N.A. 1.45, Nikon) and a Nikon DS-Qi2 camera. Images were acquired using 405(20)nm and 561(10)nm laser excitation for blue (Hoechst) and red (NileRed) dyes respectively. The cells were grown overnight in M9 medium supplemented with 1mM thiamine, 0.4% glucose and 25μg/mL kanamycin, diluted 1:1000 in fresh media and grown to early exponential phase in the absence or presence of the inducer aTc. After addition of both dyes (Hoechst at 10ug/mL and NileRed at 1ug/mL), cells were shaken at room temperature for one hour and imaged in drops of the respective growth media. Images were deconvolved using Huygens Professional (version 4.5) and further analyzed using ImageJ.

### Statistical analysis

We analyzed differences in growth rates between cultures using one-way ANOVA followed by pairwise comparisons using Tukey’s HSD to account for multiple testing (Python scipy.stats, functions f_oneway and tukey_hsd).

Spearman correlation was calculated for the fit between model predictions of binding strength and the number of obtained ChIP-sequencing reads per 1000bp window using Matlab R2019b. P-values for WT and basepair bias energy matrix predictions were P < 0.001 and for the average over random matrices P < 0.01. Peak calling on ChIP-sequencing data is described under ‘ChIP-sequencing analysis’.

## DATA AVAILABILITY

The datasets produced in this study are available in the Gene Expression Omnibus database and the computer code on GitHub (https://github.com/balaji-srinivasan-santhanam/peaks).

## ACKNOWLEDGEMENT

We thank M. Erhardt for the donation of strains, T. Bergmiller, R. Chait, K. Jain, M. La Fortezza and the ETH Zurich ScopeM facility for their support with fluorescence microscopy, N. Gnyliukh for help with Western blot experiments, and T. Friedlander and J. Crocker for useful discussions. CI was supported by a DOC Fellowship of the Austrian Academy of Sciences, the Wellcome Trust [225565/Z/22/Z] and an ETH Zurich Postdoctoral Fellowship (19-2 FEL-74). Library preparation and sequencing of ChIP seq samples was performed by the Next Generation Sequencing Facility at Vienna BioCenter Core Facilities (VBCF), member of the Vienna BioCenter (VBC), Austria.

## AUTHOR CONTRIBUTIONS

CI, CF, GT and CCG conceived the study together. CI designed the experiments and carried them out together with CF and BW (growth experiments) and TW (ChIP-sequencing). CI, FMP and BS analyzed the ChIP sequencing data. BW and BK performed and analyzed protein level experiments. CI wrote the initial draft of the manuscript and revised it with CCG, GT and BW.

