## Supplementary Figures for "Non-cognate molecular interactions impact bacterial growth"

**Supplementary Information for**
**Non-cognate molecular interactions impact bacterial growth**

Claudia Igler[1,2], Bryan Wu[1], Claire Fourcade[1], Bor Kavčič[1], Torsten Waldminghaus[3], Florian M. Pauler[1], Balaji Santhanam[4], Gašper Tkačik[1], Călin C. Guet[1,*]

[1] IST Austria, Am Campus 1, 3400 Klosterneuburg, Austria

[2] School of Biological Sciences, University of Manchester, Manchester, UK

[3] Centre for Synthetic Biology, Technische Universität Darmstadt, Darmstadt, Germany

[4] Department of Systems Biology, Columbia University, 1150 St. Nicholas Avenue, New York, NY 10032

[*] Corresponding author: Călin C. Guet; IST Austria, Am Campus 1, 3400 Klosterneuburg, Austria; +43 2243 9000 4001

**SUPPLEMENTARY FIGURES**

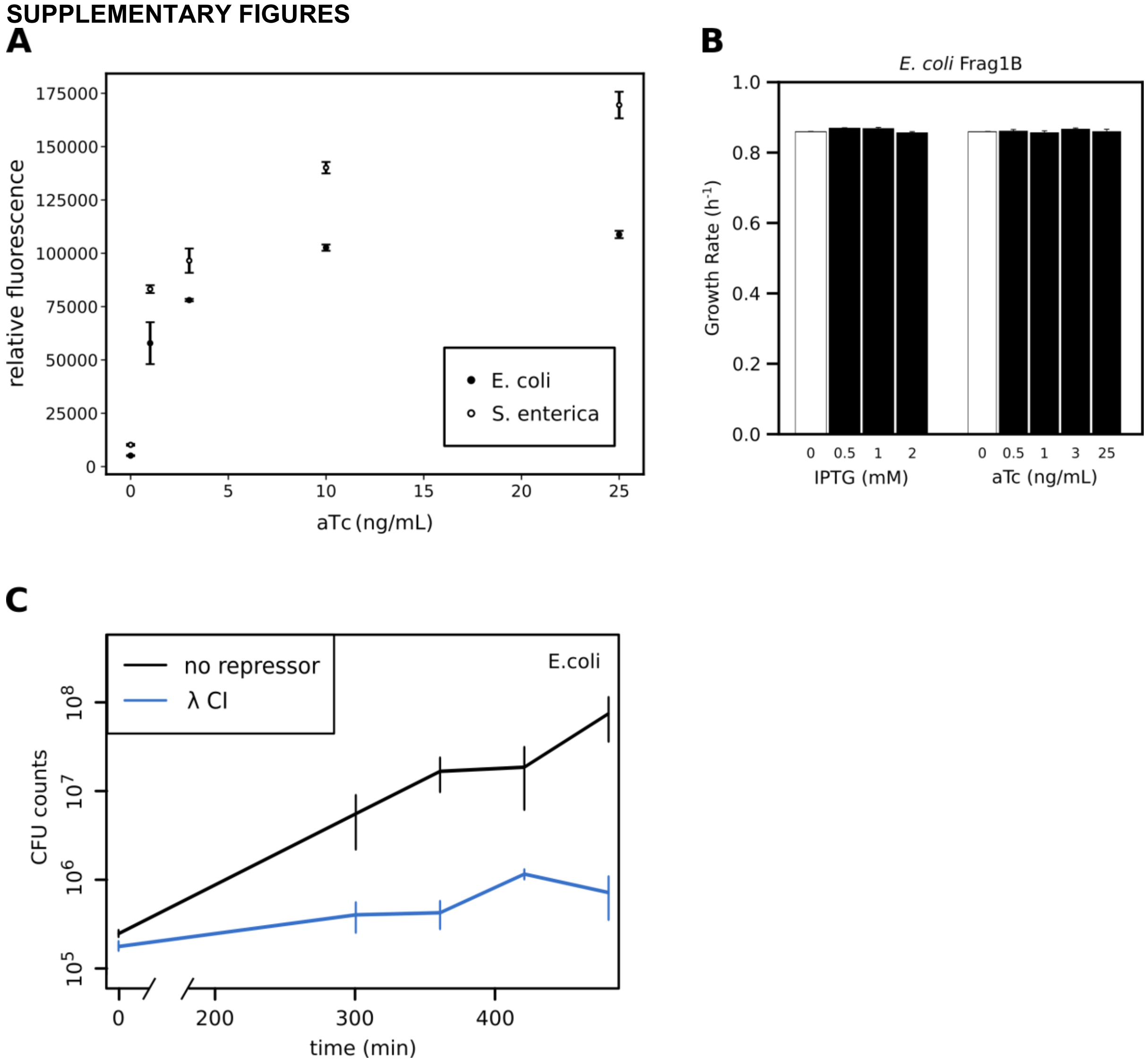


**Figure S1. Controls for growth effects.**
**(A)** *E. coli* or *S. enterica* cells containing pZS21 plasmids carrying an inducible fluorescence gene (pZS21-GFP) were grown in minimal media with glucose and induced with 0,1,3,10 or 25 ng/ml aTc. Fluorescence was measured after 11h of induction and normalized by $OD_{600}$. Error bars show standard deviation over 3 replicates. **(B)** Growth rates (per hour) for *E. coli* Frag1B cells, with constitutive LacI and TetR expression, grown in minimal media that are induced with 0, 0.5, 1 and 2mM IPTG (to relieve LacI binding), and induced with 0, 0.5, 1, 3 and 25ng/ml aTc (to relieve TetR binding). 0mM IPTG / 0ng/mL aTc shows growth in the presence of both LacI and TetR binding. **(C)** CFU counts for *E. coli* cells grown in minimal media in a shaking incubator, either expressing (25ng/mL aTc, blue) or not expressing (black) λ CI**,** sampled at 0, 300, 360, 420 and 480 min. Error bars give standard error over 3-8 replicates.

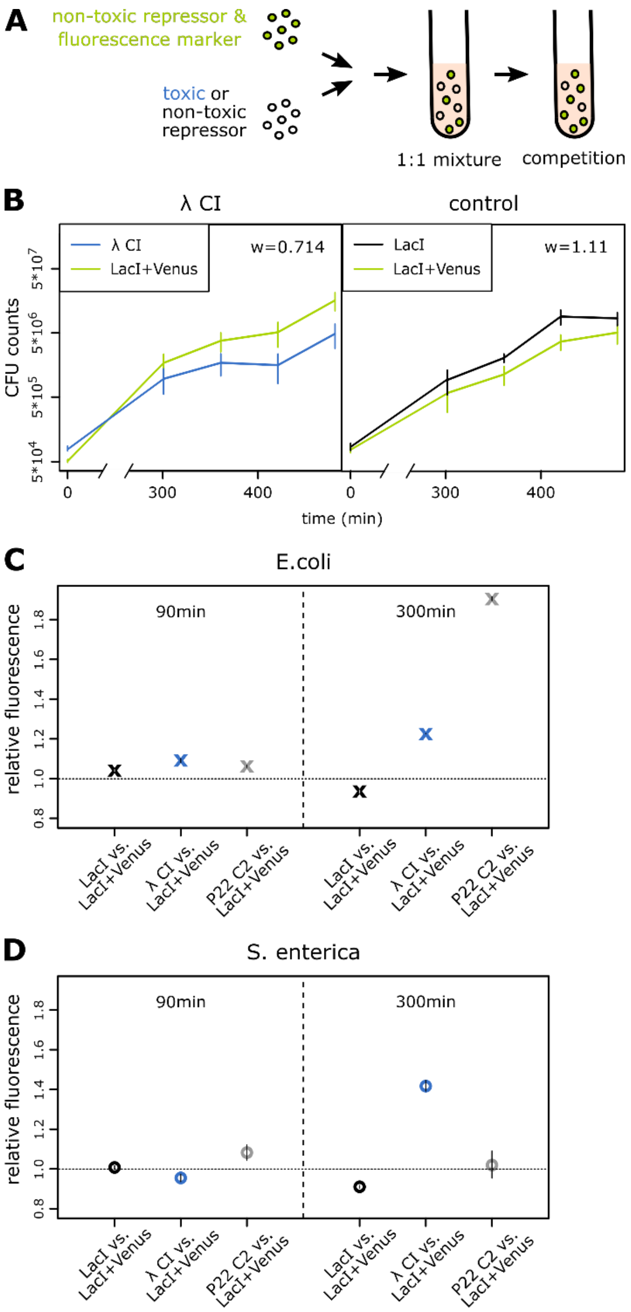


**Figure S2. Competition assays reveal the fitness cost of repressor expression.** **(A)** Cells containing plasmids with a repressor that is affecting growth when expressed (λ CI, P22 C2) or with a repressor not affecting growth when expressed (LacI) were mixed 1:1 with cells containing a plasmid with the repressor not affecting growth and a separately, constitutively expressed Venus marker (LacI+Venus). Note that LacI is also present on the chromosome of *E. coli*, where its binding has a minimal fitness cost (Fig. S1), and that additional LacI expression is not costly (Fig. 3B). Competition was performed in minimal media with glucose and **(B)** 0.8ng/mL aTc or **(C,D)** 25ng/mL aTc. **(B)** Colony forming units (CFU) for competitions between *E. coli* cells carrying LacI+Venus and *E. coli* cells carrying **(left)** λ CI or **(right)** LacI were determined at the beginning and after 300, 360, 420 and 480min. Relative fitness w was calculated as log ratio between population growth of cells carrying λ CI or LacI and that of LacI+Venus (Methods). The venus marker carried a cost as relative growth of *E. coli* without the marker is 1.11. Expression of λ CI was even more costly, as fitness of λ CI-carrying cells relative to LacI+Venus -carrying cells was w=0.74. Error bars show standard error over 4 to 14 replicates. **(C,D)** Competition was performed in minimal media with glucose and 25ng/mL aTc and fluorescence was used as a measure of the relative change in the cells carrying LacI-Venus. Relative fluorescence was calculated for **(C)** *E. coli* or **(D)** *S. enterica* between induced and non-induced samples of cell mixtures of LacI+Venus together with LacI (control), λ CI or P22 C2 after **(left)** 90min or **(right)** 300min of competition; error bars show relative errors.

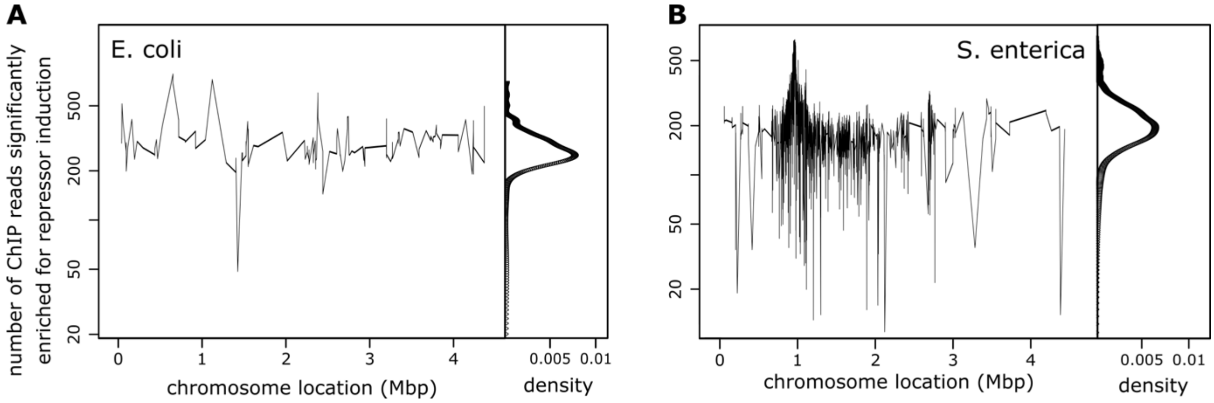


**Figure S3. ChIP-sequencing read-depth significantly enriched for repressor induction across the *E. coli* and *S. enterica* genomes.**
**(A,B)** Distributions of ChIP-sequencing reads for the regions found to be significantly enriched when using a lenient peak-calling threshold (Benjamini-Hochberg false-discovery rate of less than 30%) in the experiment with λ CI over the experiment without λ CI across the **(A)** *E.coli* or **(B)** *S. enterica* genome. On the right density plots of enriched read numbers are given. None of the apparent peaks in **(A)** or **(B)** encodes for an essential gene, nor one obviously involved in cell division.

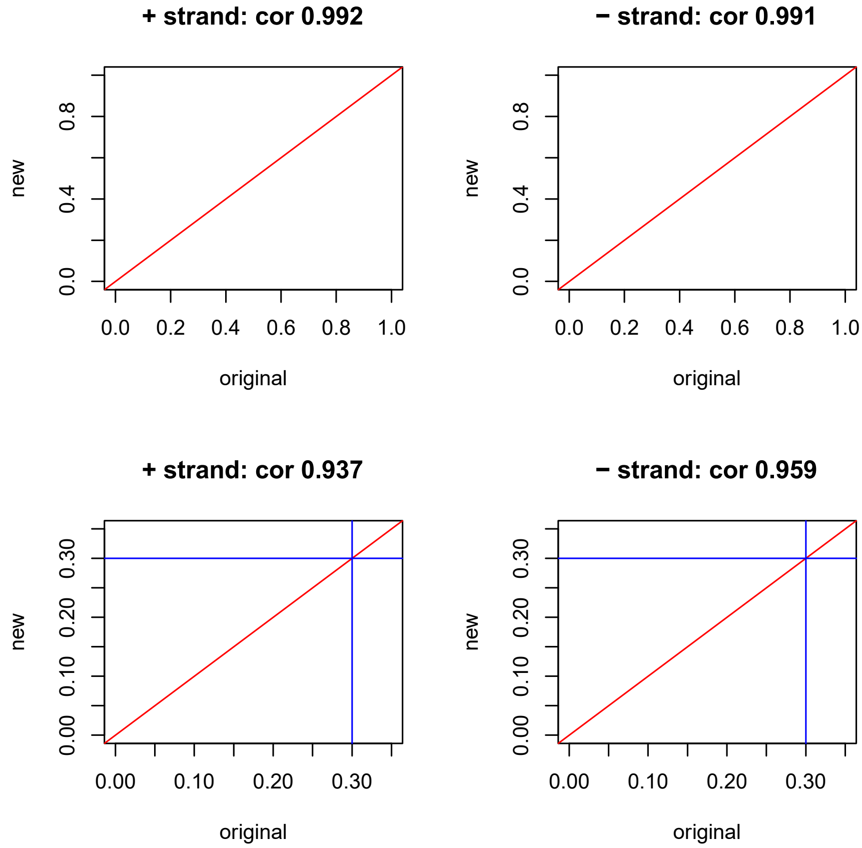


**Fig. S4. Correlation between different FDR calculation approaches.**
FDR calculation was compared between using the induced *E. coli* – λ CI sample only (original) and using both induced (ChIP) and uninduced (control) samples (new) by randomly partitioning them and assigning experiment labels to the two partitions. This process is repeated for the forward (+) and the reverse (–) strand (left and right plots respectively). Scatterplots show that FDR with random label swaps as well as FDR calculated using our original method are largely concordant (correlation > 0.9).

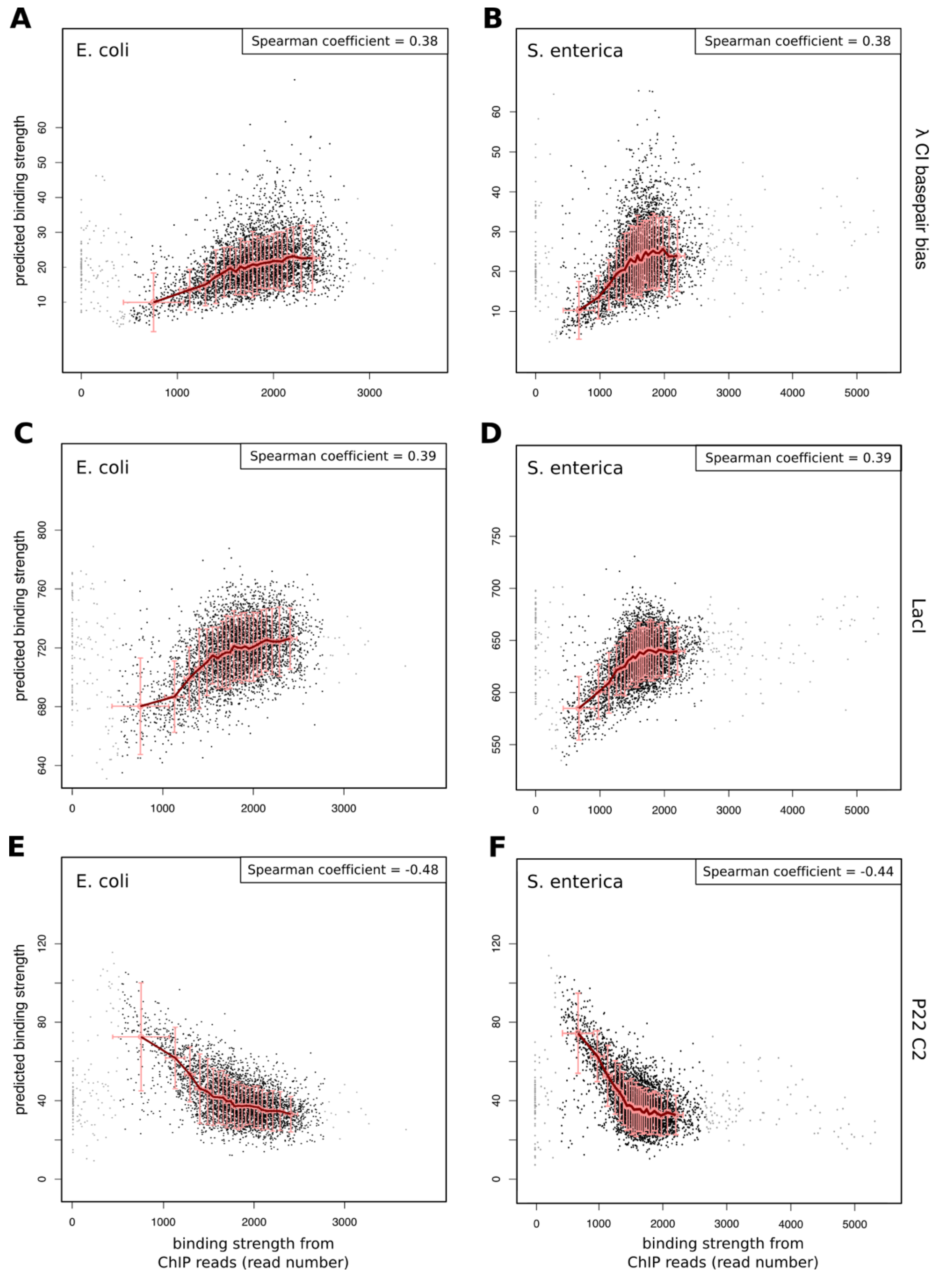


**Fig. S5. Fit of λ CI basepair bias, LacI WT and P22 C2 WT energy predictions with the λ CI ChIP-sequencing data.**

Fit between binding strength predictions of a simple thermodynamic model using an energy matrix that conserves **(A,B)** only the λ CI basepair bias or the WT energy matrices for **(C,D)** LacI and **(E,F)** P22 C2 binding (together with the corresponding WT offsets) and the λ CI ChIP-sequencing reads across 1000bp windows along the **(A,C,E)** *E. coli* or **(B,D,F)** *S. enterica* genome. The Spearman coefficients calculated for the correlations are shown. The magnitudes of the Spearman correlations for LacI and P22 C2 predictions are similar to the ones with the λ CI basepair bias, but negative (inverse) for P22 C2. As LacI has a similar basepair bias as λ CI, but P22 C2 an inverse one, this further supports that DNA sequences with the correct basepair bias provide enough recognition motif for λ CI to bind with low affinity. Note that Spearman correlation only considers the qualitative fit, but the range of predicted binding strengths for λ CI energy predictions is substantially larger than for either LacI or P22 C2.

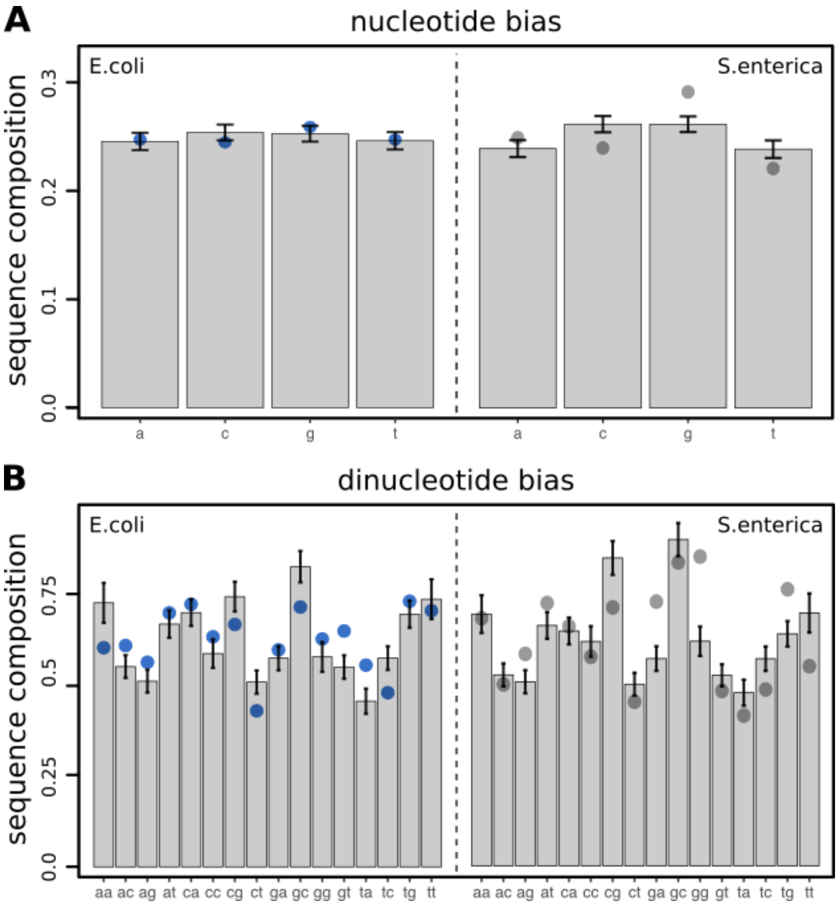


**Fig. S6. Nucleotide bias in ChIP-sequencing data.**
Bar plots show mean and standard deviation of the background sequence composition (i.e. for regions that were not enriched in ChIP-sequencing data) as compared to the sequence composition in enriched regions (blue and grey dots for *E. coli* and *S. enterica* respectively).

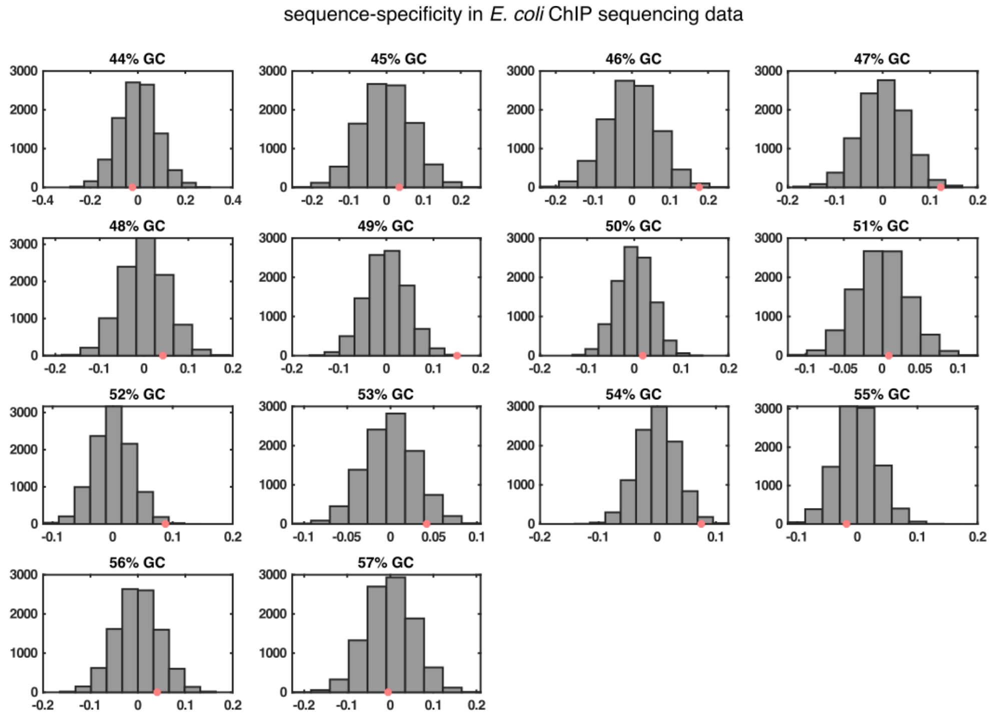


**Fig. S7. Sequence-dependence of λ CI binding to the *E. coli* genome.**
1000bp fragments of the *E. coli* genome were divided into bins of equal GC percentage (ranging from 44-57%). Histograms show the difference in Spearman correlation between the wildtype energy matrix and the one that conserves basepair bias for 10000 random assignments of ChIP reads to genome regions, whereas red dots show the difference for the real ChIP read assignment. We only included bins of a certain GC composition when they contained at least 100 genome fragments.

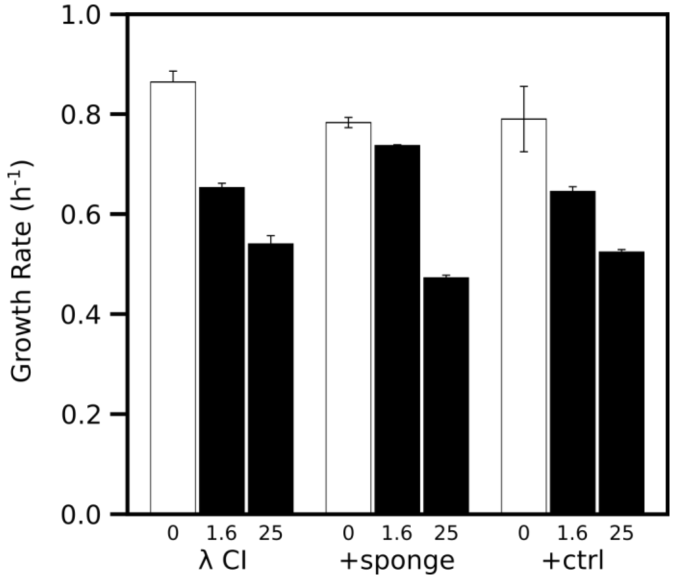


**Figure S8. Titration of λ CI by target binding sites on a high-copy plasmid.** Growth rates (per hour) are shown for *E. coli* cells containing the low-copy plasmid carrying λ CI or containing the λ CI plasmid together with a high-copy plasmid with either four natural λ CI binding sites (+sponge) or the four sites replaced with random DNA sequences (+ctrl). λ CI was induced at low (1.6ng/mL aTc induction) or high (25ng/mL aTc induction) concentrations. Uninduced growth rates are shown as white bars. Error bars show standard error over 3 replicates.

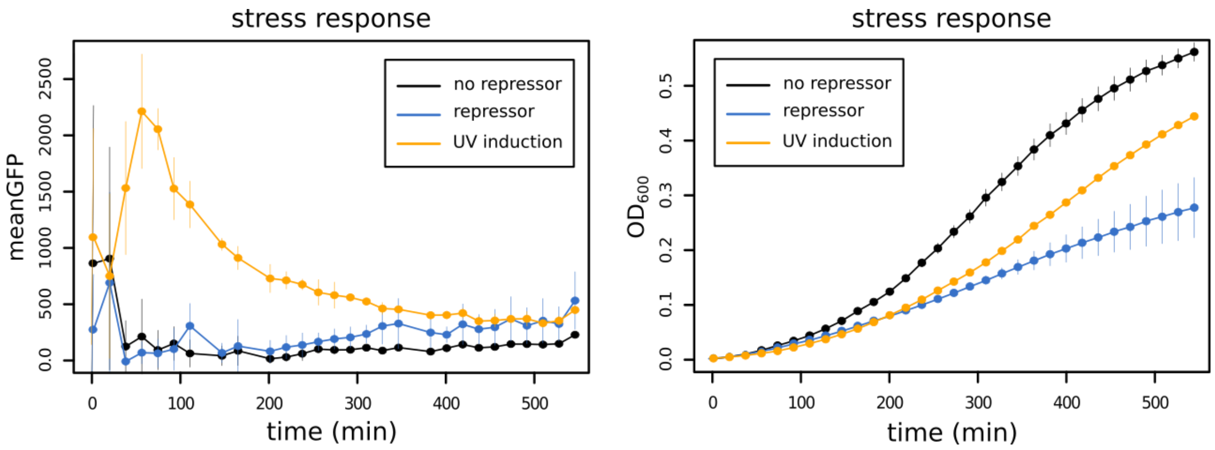


**Figure S9. Stress response induction in *E. coli* with λ CI.**
*E. coli* cells containing a stress response reporter gene ($P_{sulA}$-yfp) and λ CI were grown in minimal media with glucose. Cells were either induced for a certain stressor (25ng/mL aTc for λ CI expression (blue), or UV exposure for 30sec (yellow)), or not induced (black) and growth ($OD_{600}$) was measured over several hours. UV induction showed a spike in the stress response (mean absolute gfp fluorescence) and an initial decrease in growth ($OD_{600}$), whereas λ CI only led to a much lower stress response induction at later time points and showed an increasing growth reduction. Error bars show standard error over 3 replicates.

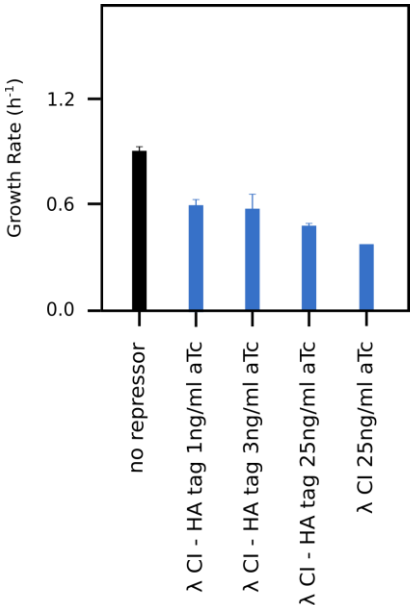


**Figure S10. HA-tagged λ CI produces the same growth patterns as wildtype repressor.** Growth rates (per minute) for *E. coli* in minimal media without (black) or with induction of λ CI-HA using 1ng/mL, 3ng/ml or 25ng/mL aTc (blue). For comparison induction of λ CI using 25ng/ml aTc in is shown as well. Error bars show standard deviation over 3 replicates.
